# Evolution and genetic architecture of disassortative mating at a locus under heterozygote advantage

**DOI:** 10.1101/616409

**Authors:** Ludovic Maisonneuve, Mathieu Chouteau, Mathieu Joron, Violaine Llaurens

**Affiliations:** Institut de Systematique, Evolution, Biodiversité (UMR7205), Museum National d’Histoire Naturelle, CNRS, Sorbonne-Université, EPHE, Université des Antilles – CP50, 57 rue Cuvier, 75005 Paris, France; Laboratoire Ecologie, Evolution, Interactions Des Systèmes Amazoniens (LEEISA), USR 3456, Université De Guyane, IFREMER, CNRS Guyane – 275 route de Montabo, 97334 Cayenne, French Guiana; (CEFE), Univ Montpellier, CNRS, EPHE, IRD, Univ Paul Valéry Montpellier 3 – Montpellier, France

**Keywords:** Disassortative mating, Supergene, Frequency dependent selection, genetic load, mate preference, *Heliconius numata*

## Abstract

The evolution of mate preferences may depend on natural selection acting on the mating cues and on the underlying genetic architecture. While the evolution of assortative mating with respect to locally adapted traits has been well-characterized, the evolution of disassortative mating is poorly characterized. Here we aim at understanding the evolution of disassortative mating for traits under strong balancing selection, by focusing on polymorphic mimicry as an illustrative example. Positive frequency-dependent selection exerted by predators generates local selection on wing patterns acting against rare variants and promoting local monomorphism. This acts across species boundaries, favouring Mullerian mimicry among defended species. In this well-characterized adaptive landscape, polymorphic mimicry is rare but is observed in a butterfly species, associated with polymorphic chromosomal inversions. Because inversions are often associated with recessive deleterious mutations, we hypothesize they may induce heterozygote advantage at the color pattern locus, putatively favoring the evolution of disassortative mating. To explore the conditions underlying the emergence of disassortative mating, we modeled both a trait locus (colour pattern for instance), subject to mutational load, and a preference locus. We confirm that heterozygote advantage favors the evolution of disassortative mating and show that disassortative mating is more likely to emerge if at least one allele at the trait locus is free from any recessive deleterious mutations. We modelled different possible genetic architectures underlying mate choice behaviour, such as self referencing alleles, or specific preference or rejection alleles. Our results showed that self referencing or rejection alleles linked to the color pattern locus can be under positive selection and enable the emergence of disassortative mating. However rejection alleles allow the emergence of disassortative mating only when the color pattern and preference loci are tightly linked. Our results therefore provide relevant predictions on both the selection regimes and the genetic architecture favoring the emergence of disassortative mating and a theoretical framework in which to interprete empirical data on mate preferences in wild populations.

## Introduction

Mate preferences often play an important role in shaping trait diversity in natural populations, but the mechanisms responsible for their emergence often remain to be characterized. While the evolution of assortative mating on locally adapted traits is relatively well understood (Cara et al., 2008; Otto et al., 2008; Thibert-Plante and Gavrilets, 2013), the selective forces involved in the evolution of disassortative mating are still largely unknown. Disassortative mating, *i.e*. the preferential mating between individuals displaying different phenotypes, is a rare form of mate preference (Jiang et al., 2013). In populations where individuals tend to mate with phenotypically distinct partners, individuals with a rare phenotype have a larger number of available mates, resulting in a higher reproductive success. By generating negative frequency-dependent selection on mating cues, disassortative mating is often regarded as a process generating and/or maintaining polymorphism within populations. Obligate dis-assortative mating leads to the persistence of intermediate frequencies of sexes or mating types (Wright, 1939), and promotes polymorphism (e.g. the extreme case of some Basidiomycete fungi where thousands of mating types are maintained (Casselton, 2002)). Disas-sortative mating can be based on different traits. Disassortative mating based on odors is known to operate in mice (Penn and Potts, 1999) and humans (Wedekind, Seebeck, et al., 1995). Odor profiles are associated with genotype at the MHC loci affecting the immune response, known to be under strong balancing selection (Piertney and Oliver, 2006). Balancing selection on MHC alleles partly stems from heterozygous advantage, whereby heterozygous genotypes might confer an ability to recognize a larger range of pathogens. Such heterozygote advantage may promote the evolution of disassortative mating (Tregenza and Wedell, 2000). Extreme examples of heterozygote advantage are observed for loci with reduced homozygote survival. In the seaweed fly *Coelopa frigida* heterozygotes (*αβ*) at the locus Adh have a higher fitness than homozygotes (*αα* or *ββ*) (RK Butlin et al., 1984; Mérot et al., 2019) and females prefer males with a genotype that differs from their own (Day and Butlin, 1987). In the white-throated sparrow *Zonotrichia albicollis*, strong disassortative mating is known to operate with respect to the color of the head stripe and associated with chromosomal dimorphism (Throneycroft, 1975). This plumage dimorphism is associated with a spectacular chromosomal polymorphism (Tuttle et al., 2016), with a complete lack of homozygous individuals for the rearranged chromosome (Horton et al., 2013).

While the fitness advantage of disassortative mating targeting loci with overdominance seems straightforward, the genetic basis of disassortative preferences remains largely unknown. One exception is the self-incompatibility system in *Brassicaceae* where the S-locus determines a specific rejection of incompatible pollens (Hiscock and McInnis, 2003). S-haplotypes contain tightly linked, co-evolved SCR and SRK alleles, encoding for a protein of the pollen coat and a receptor kinase located in the pistil membrane respectively, preventing fertilization from self-incompatible pollen due to specific receptor-ligand interactions. Self-rejection has also been proposed as an explanation for the disassortative mating associated with odor in humans. Body odors are strongly influenced by genotypes at the immune genes HLA and rejection of potential partners has been shown to be related to the level of HLA similarity, rather than to a particular HLA genotype (Wedekind and Füri, 1997). In the white-throated sparrow, disassortative mating results from specific preferences for color plumage that differ between males and females; *tan*-striped males are preferred by all females while *white*-striped females are preferred by all males (Houtman and Falls, 1994). Different mechanisms leading to mate preferences and associated genetic architecture can be hypothesized, that may involve the phenotype of the chooser. Based on the categories described by Kopp et al., 2018, we assume that disassortative mating can emerge from two main mechanisms. (1) *Self-referencing*, when an individual uses its own signal to choose its mate, which may generate a disassortative mating that depends on the phenotypes of both the choosing and the chosen partners. (2) Preferences for or rejection of a given phenotype in the available partners (*recognition/trait* hypothesis), independently from the phenotype of the choosing partner, may also enable the emergence of disassortative mate preferences. These two mechanisms could involve a two locus architecture where one locus controls the mating cue and the other one the preference towards the different cues (Kopp et al., 2018). The level of linkage disequilibrium between the two loci could have a strong impact on the evolution of disassortative mating. In models investigating the evolution of assortative mating on locally-adapted traits, theoretical simulations have demonstrated that assortative mating is favored when the preference and the cue loci are linked (Kopp et al., 2018).

Here we explore the evolutionary forces leading to the emergence of disassortative mating. We use as a model system the specific case of the butterfly species *Heliconius numata*, where high polymorphism in wing pattern is maintained within populations (Joron, Wynne, et al., 1999) and strong disassortative mating operates between wing pattern forms (Chouteau, Llaurens, et al., 2017). *H. numata* butterflies are chemically-defended (Arias et al., 2016; Chouteau, Dezeure, et al., 2019), and their wing patterns act as warning signals against predators (Chouteau, Arias, et al., 2016a). A a local scale, natural selection on local mimicry usually leads to the fixation of a single warning signal shared by multiple defended species (Müllerian mimicry) (Mallet and Barton, 1989). However, local polymorphism of mimetic color patterns is maintained in certain species for instance under a balance between migration and local selection on mimicry (Joron and Iwasa, 2005). Yet, the level of polymorphism observed within populations of *H. numata* (Joron, Wynne, et al., 1999) would require that the strong local selection is balanced by a very high migration rate. However, disassortative mating based on wing pattern operates in *H. numata*, with females rejecting males displaying the same color pattern (Chouteau, Llaurens, et al., 2017). Such disassortative mating could enhance local polymorphism in color pattern within this species. Nevertheless, the mode of evolution of a disassortative mating is unclear, notably because preferences for dissimilar mates should not be favoured if natural selection by predators on adult wing pattern acts against rare morphs (Chouteau, Arias, et al., 2016b). Building on this well-documented case study, we use a theoretical approach to provide general predictions on the evolution of disassortative mating in polymorphic traits, and on expected genetic architecture underlying this behavior.

Variation in wing color pattern in *H. numata* is controlled by a single genomic region, called the supergene P (Joron, Papa, et al., 2006), displaying dictinct chromosomal inversions combinations, each associated with a distinct mimetic phenotype (Joron, Frezal, et al., 2011). These inversions have recently been shown to be associated with a significant genetic load, resulting in a strong heterozygote advantage (Jay et al., 2019). We thus investigate whether a genetic load associated with locally adaptive alleles may favor the evolution of mate preference and promote local polymorphism. We then explore two putative genetic architectures for mate preferences based on (1) *self referencing* and (2) based on a *recognition/trait* rule, and test for their respective impacts on the evolution of disassortative mating. Under both hypotheses, we assumed that the mating cue and the mating preference were controlled by two distinct loci, and investigate the effect of linkage between loci on the evolution of disassortative mating.

## Methods

### Model overview

Based on earlier models of Müllerian mimicry (Joron and Iwasa, 2005; Llaurens, Billiard, et al., 2013), we describe the evolution of mate preferences based on color pattern using ordinary differential equations (ODE). We track the density of individuals carrying different genotypes combining the alleles at the locus *P* controlling mimetic color pattern and at the locus *M* underlying sexual preference. We assume a diploid species, so that each genotype contains four alleles.

The set of all possible four-allele genotypes is defined as 𝒢 = 𝒜_*P*_ × 𝒜_*P*_ × 𝒜_*M*_ × 𝒜_*M*_ where 𝒜_*P*_, 𝒜_*M*_ are the set of alleles at locus *P* and *M* respectively. A given genotype is then an quadruplet of the form (*p*_*m*_, *p*_*f*_, *m*_*m*_, *m*_*f*_) with *p*_*m*_ ∈ 𝒜_*P*_ and *m*_*m*_ ∈ 𝒜_*M*_ (resp. *p*_*f*_ and *m*_*f*_) being the alleles at loci *P* and *M* on the maternal (resp. paternal) chromosomes. A recombination rate *ρ* between the color pattern locus *P* and the preference locus *M* is assumed.

We consider two geographic patches numbered 1 and 2 where those genotypes can occur. For all (*i, n*) ∈ 𝒢 × {1, 2} we track down the density of individuals of each genotype *i* within each patch *n, N*_*i,n*_ trough time. Following previous models, polymorphism in mimetic color pattern is maintained within each of the two patches, by a balance between (1) local selection on color pattern in opposite directions in the two patches and (2) migration between patches.

The evolution of genotype densities through time, for each patch, is influenced by predation, mortality, migration between patches and reproduction, following the general equations :

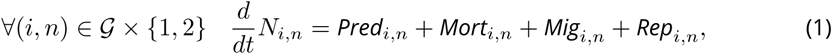

where *Pred*_*i,n*_, *Rep*_*i,n*_, *Mig*_*i,n*_, and *Mort*_*i,n*_ described the respective contributions of these four processes to the change in density of genotype *i* within each patch *n*. The computation of each of these four contributions is detailed in specific sections below. All variables and parameters are summarized in Table 1 and 2 respectively.

**Table 1.**
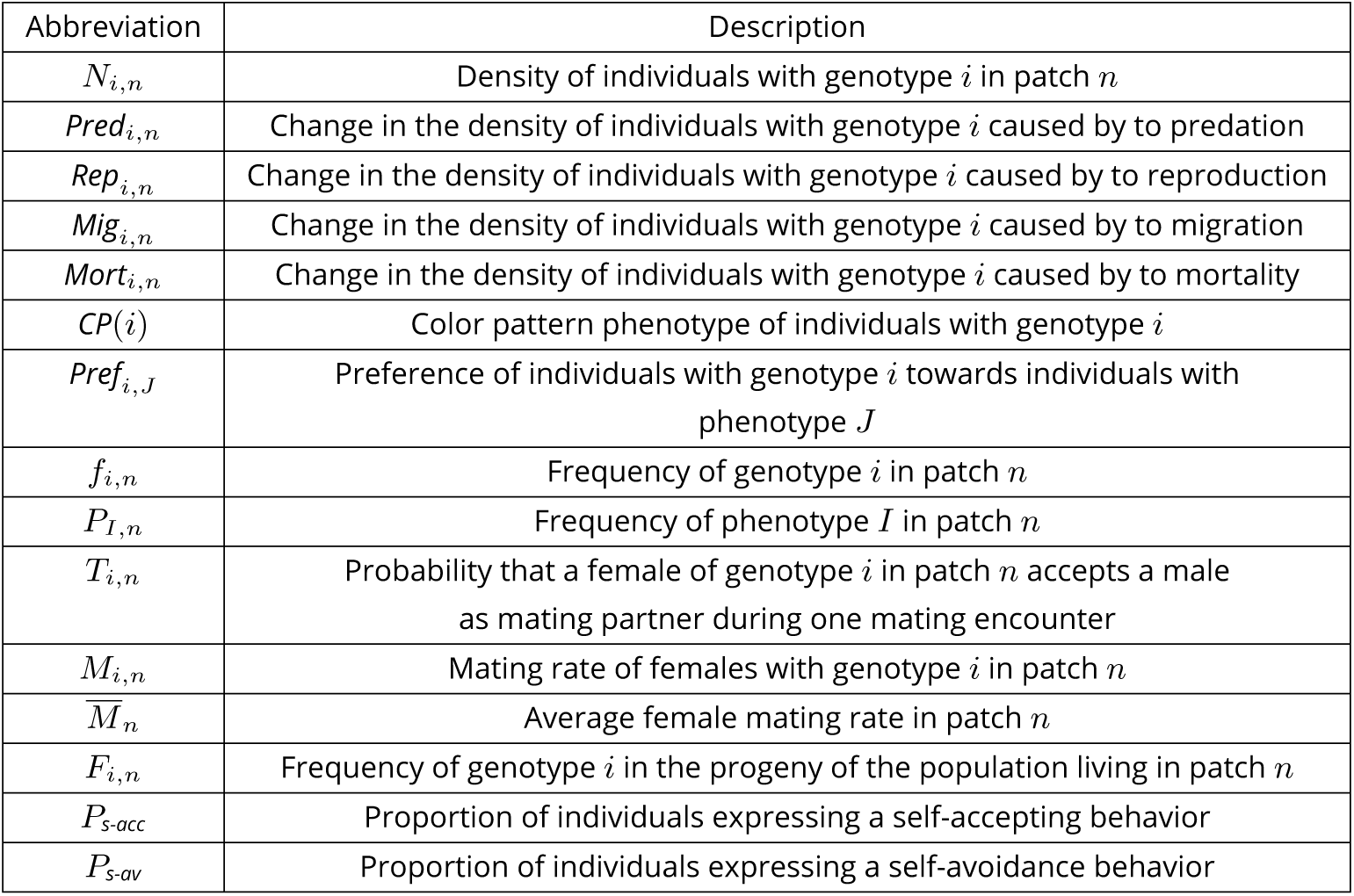
Description of variables used in the model.

**Table 2.**
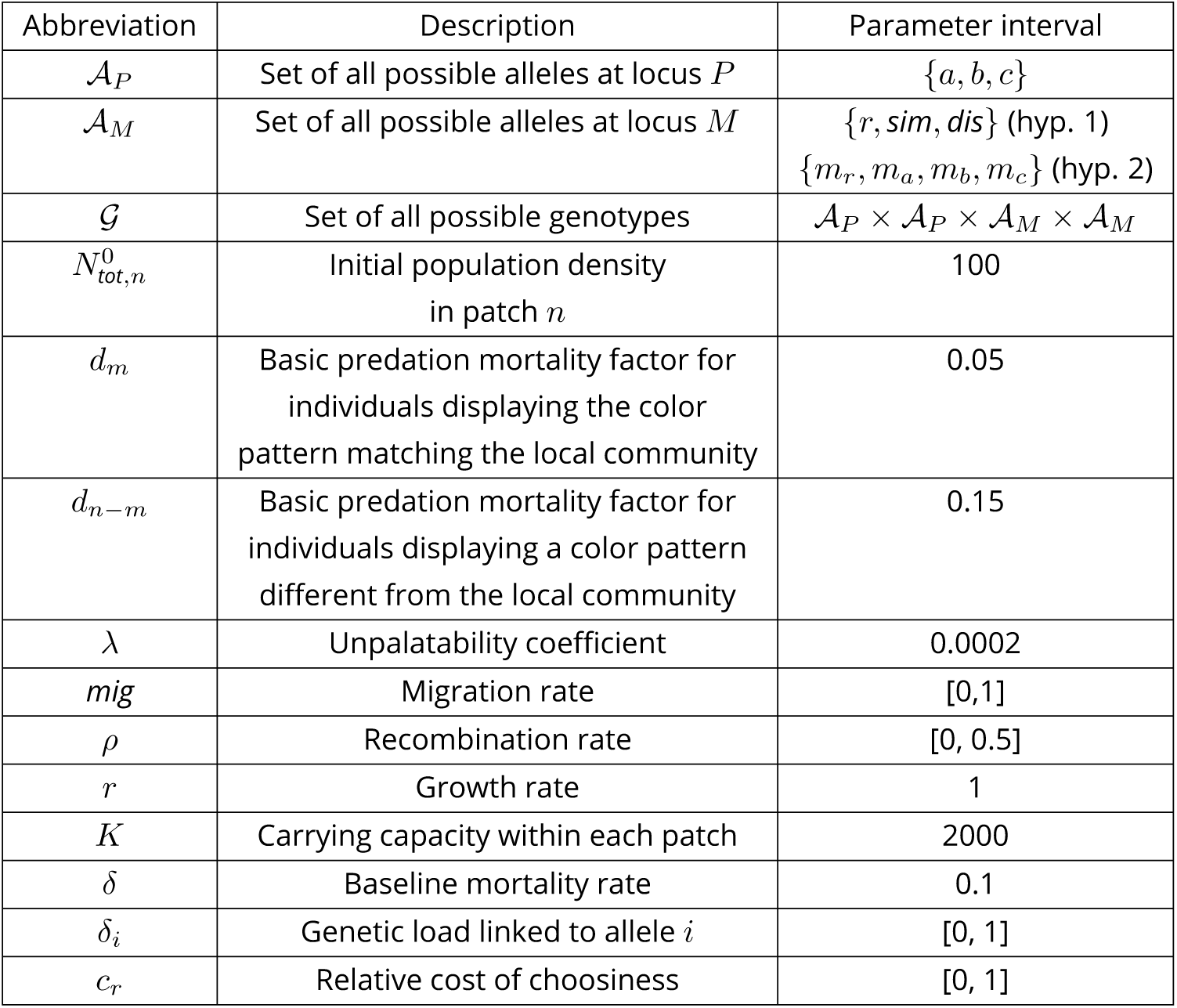
Description of parameters used in the model and range explored in simulations.

| Abbreviation | Description | Parameter interval |
| --- | --- | --- |
| $\mathcal{A}_P$ | Set of all possible alleles at locus $P$ | $\{a, b, c\}$ |
| $\mathcal{A}_M$ | Set of all possible alleles at locus $M$ | $\{r, sim, dis\}$ (hyp. 1)<br>$\{m_r, m_a, m_b, m_c\}$ (hyp. 2) |
| $\mathcal{G}$ | Set of all possible genotypes | $\mathcal{A}_P \times \mathcal{A}_P \times \mathcal{A}_M \times \mathcal{A}_M$ |
| $N_{tot,n}^0$ | Initial population density in patch $n$ | 100 |
| $d_m$ | Basic predation mortality factor for individuals displaying the color pattern matching the local community | 0.05 |
| $d_{n-m}$ | Basic predation mortality factor for individuals displaying a color pattern different from the local community | 0.15 |
| $\lambda$ | Unpalatability coefficient | 0.0002 |
| $mig$ | Migration rate | [0,1] |
| $\rho$ | Recombination rate | [0, 0.5] |
| $r$ | Growth rate | 1 |
| $K$ | Carrying capacity within each patch | 2000 |
| $\delta$ | Baseline mortality rate | 0.1 |
| $\delta_i$ | Genetic load linked to allele $i$ | [0, 1] |
| $c_r$ | Relative cost of choosiness | [0, 1] |

Since our ODE model describes the change in genotype densities at a population level, this amounts to considering that predation, migration, reproduction and survival occur simultaneously (see Equation (1)). In a large population, we can assume that predation, migration, reproduction and survival indeed occur in different individuals at the same time. Such a model implies that generations are overlapping and that there is no explicit ontogenic development: each newborn individual instantaneously behaves as an adult individual and can immediately migrate and reproduce. Our deterministic model provides general predictions while ignoring the effects of stochastic processes such as genetic drift.

#### Mimetic color pattern alleles at locus *P*

At the color pattern locus *P*, three alleles are assumed to segregate, namely alleles *a, b* and *c*, encoding for phenotypes *A, B* and *C* respectively. The set of alleles at locus *P* is then 𝒜_*P*_ = {*a, b, c*}. We assume strict dominance among the three alleles with *a > b > c* in agreement with the strict dominance observed among supergene *P* alleles within natural populations of *H. numata* (Le Poul et al., 2014) and in other supergenes (Küpper et al., 2016; Tuttle et al., 2016; Wang et al., 2013). The three color pattern phenotypes are assumed to be perceived as categorically different by both mating partners and predators. We note *CP* the function translating each genotype *i* into the corresponding color pattern phenotype 𝒢. For example, for all (*m*_*m*_, *m*_*f*_) ∈ 𝒜_*M*_ ×𝒜_*M*_, *CP*((*a, b, m*_*m*_, *m*_*f*_)) = *A* because allele *a* in dominant over *b* and the color pattern phenotype depends only on alleles at locus *P*. Each color pattern allele is also assumed to carry an individual genetic load expressed when homozygous.

#### Preference alleles at locus *P*

We investigate the evolution of mate preference associated with color patterns, exploring in particular the conditions enabling the evolution of disassortative mating. We assume a single choosy sex: only females can express preferences toward male phenotypes, while males have no preference and can mate with any accepting females. Female preferences toward males displaying different color patterns are controlled by the locus *M*. We assume two different models of genetic architecture underlying mate preferences: alleles at locus *M* determine either (1) a preference toward similar or dissimilar phenotypes, which therefore also depends on the phenotype of the choosing individual, following the *self-referencing* hypothesis or (2) a preference toward a given color pattern displayed by the mating partner, independent of the color pattern of the choosing individual, following the *recognition/trait* hypothesis.

### Predation

The probability of predation on individuals depends on their mimetic color patterns controlled by the locus *P*. Predation is determined in our model by a basic (patch-specific) effect of the local community of prey favouring one of the wing patterns locally (local adaptation through mimicry), itself modulated by positive frequency dependence of the different wing patterns controlled by P, within the focal species population. This is detailed below.

#### Divergent local adaptation in color pattern

Local selection exerted by predators promotes convergent evolution of wing color patterns among defended species (*i.e*. Müllerian mimicry, (Müller, 1879)), forming so-called mimicry rings composed of individuals from different species displaying the same warning signal within a locality. Mimicry toward the local community of defended prey therefore generates strong local selection on color pattern and the direction of this selection then varies across localities (Sherratt, 2006).

Here we assume two separate populations exchanging migrants of an unpalatable species involved in Müllerian mimicry with other chemically-defended species. Local communities of species involved in mimicry (*i.e*. mimicry rings) differ across localities. We consider two patches occupied by different mimetic communities: population 1 is located in a patch where the local community (*i.e*. other chemically-defended species, not including *H. numata*) mostly displays phenotype A, and population 2 in a patch where the mimetic community mostly displays phenotype B. This spatial variation in mimicry rings therefore generates a divergent selection favouring distinct locally adapted phenotypes. Note that allele *c*, and corresponding phenotype *C* is non-mimetic in both patches and at a disadvantage in both patches. Every individual of the focal (polymorphic) species is exposed to a predation risk modulated by its resemblance to the local mimetic community of butterflies. Each genotype *i* in population *n* (with (*i, n*) ∈ 𝒢 × {1, 2}) suffers from a basic predation mortality factor *d*_*i,n*_. This parameter is lower for individuals displaying the phenotype mimetic to the local community (*i.e*. the phenotype *A* in population 1 and *B* in population 2). Individuals displaying phenotype *C* being non-mimetic in both patches, suffer from a high predation risk in both patches.

Here, to simplify, we consider that this basic mortality factor takes the value *d*_*m*_ for the locally mimetic phenotype (*A* in patch 1, *B* in patch 2), and *d*_*n-m*_ for the locally non-mimetic phenotypes (*B* and *C* in patch 1, *A* and *C* in patch 2). We therefore introduce parameters *d*_*n-m*_ and *d*_*m*_, with *d*_*n-m*_ *> d*_*m*_, as follows: the basic predation mortality factors for individuals not displaying and displaying the same color pattern as the local community respectively. For *i* ∈ 𝒢, the basic predation mortality factors of individuals with genotype *i* in patch 1 and 2 are

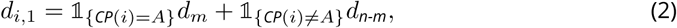

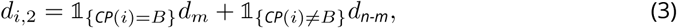

where 𝟙 is the indicator function which return 1 if the condition under brace is true and 0 else.

#### Local positive frequency-dependent predation

Predation exerted on a given phenotype depends on its match to the local mimetic environment (described by the parameter *d*_*i,n*_ for all (*i, n*) ∈ 𝒢 ×{1, 2}, see previous paragraph), but also on its own abundance in the patch as predators learn to associate warning patterns with chemical defense. This learning behavior generates positive frequency-dependent selection on color patterns (Chouteau, Arias, et al., 2016b): displaying a widely shared color pattern decreases the risk of encountering a naive predator (Sherratt, 2006). Number-dependent predator avoidance in the focal species is assumed to depend on its unpalatability coefficient (*λ*) and on the density of each phenotype within the population: the protection gained by phenotypic resemblance is greater for higher values of the unpalatability coefficient *λ*. For (*i, n*) ∈ 𝒢 × {1, 2}, the change in the density of a genotype *i* in patch *n* due to predation thus takes into account both the spatial variation in mimetic communities (using *d*_*i,n*_) modulated by the local frequency-dependent selection, and is thus described by the equation:

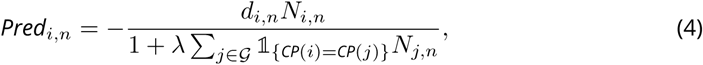

where 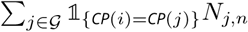 is the total density, within patch *n*, of individuals sharing the same color pattern as individuals of genotype *i*.

### Mortality

We assume a baseline mortality rate *δ*. The recessive genetic loads *δ*_*a*_, *δ*_*b*_, *δ*_*c*_ associated with the respective alleles *a, b* and *c* limit the survival probabilities of homozygous genotypes at locus *P*.

For *i* = (*p*_*m*_, *p*_*f*_, *m*_*m*_, *m*_*f*_) ∈ 𝒢, *n* ∈ {1, 2} the change in density of individuals with genotype *i* in patch *n* is given by

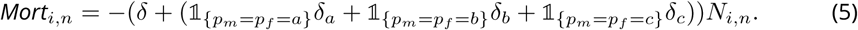

### Migration

We assume a constant symmetrical migration rate *mig* corresponding to a proportion of individuals migrating from one patch to the other, as classically assumed in population genetics models (see for instance Holt, 1985; Joron and Iwasa, 2005; Kuang and Takeuchi, 1994). The number of individuals of each of the genotypes migrating to the other patch is therefore directly proportional to their density in their source population. For (*i, n, n*′) ∈ 𝒢 × {1, 2} × {1, 2}, *n ≡ n*′, the change in the density of individuals with genotype *i* in patch *n* due to migration between patches *n* and *n*′ is given by the difference between the density of individuals coming into the patch 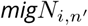 and those leaving the patch *migN*_*i,n*_:

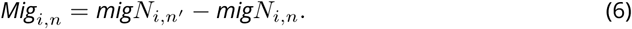

where *mig* is the migration coefficient *mig* ∈ [0, 1].

### Reproduction

In the model, the reproduction term takes into account the basic demographic parameter, the effect of mate preference controlled by locus *M* and the fecundity limitations associated with choosiness.

#### Local demography

We assume that the populations from both patches have identical carrying capacity *K* and growth rate *r*. We name *N*_*tot,n*_ the total density in patch *n*. The change in the total density due to reproduction is given by the logistic regulation function 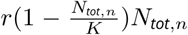. Thus for (*i, n*) ∈ 𝒢 × {1, 2}, the change in the density of genotype *i* in patch *n* generated by sexual reproduction is given by:

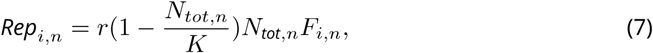

where (*F*_*i,n*_)_*i*∈𝒢_ are the frequencies of each genotype in the progeny. These frequencies depend on the behavior of the female, controlled by the preference locus *M* and on the availability of the preferred partners in the population, as detailed in the following section.

#### Mate preferences

During sexual reproduction, we assume that only one out of the two sexes expresses a mate preference, as often observed in sexual reproduction where females are usually choosier. Thus we assume females to be the choosy sex. The mate preference of female is then considered strict, implying that choosy individuals never mate with individuals displaying their non-preferred phenotype. Two hypothetical mate preference mechanisms are investigated.

Under the *self-referencing* hypothesis (hyp 1), three alleles are assumed at loci *M*, coding for (i) random mating *r*, (ii) assortative mating *sim* and (iii) disassortative *dis* respectively (see fig. S5 for more details, (𝒜_*M*_ = {*r, sim, dis*}). We assume that the *self-referencing* preference alleles *sim* and *dis* are dominant to the random mating allele *r* (see fig. S5 for more details). The dominance relationship between the *sim* and *dis* alleles is not specified however, because we never introduce these two alleles together. Note that under the *self-referencing* hypothesis (hyp. 1), mate choice depends not only on the color pattern of the male, but also on the phenotype of the female expressing the preference.

The alternative mechanism of mate preference investigated, assumes a specific recognition of color patterns acting as mating cue (*recognition/trait*, hyp. 2). Under hyp. 2, four alleles segregate at locus *M*: allele *m*_*r*_, coding for an absence of color pattern recognition (leading to random mating behavior), and *m*_*a*_, *m*_*b*_ and *m*_*c*_ coding for specific recognition of color pattern phenotypes *A, B* and *C* (𝒜_*M*_ = {*m*_*r*_, *m*_*a*_, *m*_*b*_, *m*_*c*_}). The *no preference* allele *m*_*r*_ is recessive to all the preference alleles *m*_*a*_, *m*_*b*_ and *m*_*c*_, and preference alleles are co-dominant, so that females with heterozygous genotype at locus *M* may recognize two different color pattern phenotypes. Then, the recognition enabled by preference alleles *m*_*a*_, *m*_*b*_ and *m*_*c*_ triggers either *attraction* (hyp. 2.a) or *rejection* (hyp. 2.b) toward the recognized color pattern, leading to assortative or disassortative mating depending on the genotype *i* of the female and the color pattern phenotype of the male (see figure S6 and S7 for more details).

#### Genotype frequencies in the progeny

We assume separate sexes and obligate sexual reproduction, and therefore compute explicitly the Mendelian segregation of alleles during reproduction, assuming a recombination rate *ρ* between the color pattern locus *P* and the preference locus *M*. We assume that the frequency of male and female of a given phenotype is the same. For (*i, n*) ∈ 𝒢 × {1, 2}, the frequency of genotype *i* in the progeny in patch *n* (*F*_*i,n*_) then also depends on the frequencies of each genotype in the patch and on the mate preferences of females computed in equation (13). We introduce the preference coefficients (*Pref*_*i,J*_)_(*i,J*)∈ 𝒢×{*A,B,C*}_. These coefficients depend on the alleles at locus *M* as detailed in the next section. For (*i, J*) ∈ 𝒢 × {*A, B, C*} the preference coefficient *Pref*_*i,J*_ is defined as *Pref*_*i,J*_ = 1 when females with genotype *i* accept males with phenotype *J* as mating partners and *Pref*_*i,J*_ = 0 otherwise.

For *i* ∈ 𝒢, *n* ∈ {1, 2}, we define *T*_*i,n*_ as the probability that a female of genotype *i* in patch *n* accepts a male during a mating encounter (see (Otto et al., 2008)):

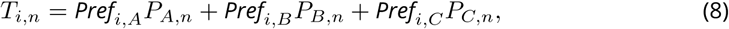

where for 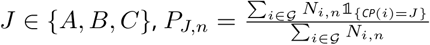 denotes the frequency of phenotype *J* in patch *n*.

Because choosy individuals might have a reduced reproductive success due to limited mate availability (Kirkpatrick and Nuismer, 2004; Otto et al., 2008), we also assume a relative fitness cost associated with choosiness. This cost is modulated by the parameter *c*_*r*_. When this cost is absent (*c*_*r*_ = 0), females have access to a large quantity of potential mates, so that their mating rate is not limited when they become choosy (“Animal” model). When this cost is high (*c*_*r*_ = 1), females have access to a limited density of potential mates, so that their mating rate tends to decrease when they become choosy (“Plant” model). Intermediate values of *c*_*r*_ imply that females can partially recover the fitness loss due to the encountering of non-preferred males towards reproduction with other males. This cost of choosiness is known to limit the evolution of assortative mating (Otto et al., 2008) and may thus also limit the emergence of disassortative mating.

Following (Otto et al., 2008) we compute the mating rate *M*_*i,n*_ of a female with genotype *I* in patch *n* :

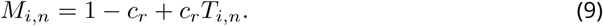

We note 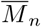 the average mating rate in patch *n* defined as

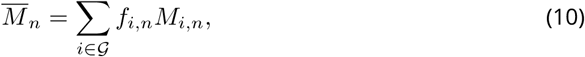

where for (*i, n*) ∈ 𝒢 × {1, 2} *f*_*i,n*_, is the frequency of genotype *i* in patch *n*.

For (*j, k*) ∈ 𝒢^2^, the quantity

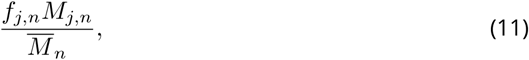

is the probability that, given that a female has mated in patch *n*, this female is of genotype j, and

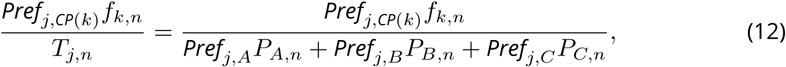

is the probability that, given that a female of genotype *j* has mated in patch *n*, its mate is a male of genotype *k*, depending on female preference and availability of males carrying genotype *k*.

For (*i, n*) ∈ 𝒢 × {1, 2}, the frequency of genotype *i* in the progeny of the population living in patch *n* is

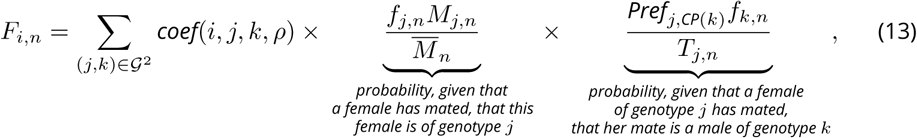

where *coef* (*i, j, k, ρ*) controls the mendelian segregation of alleles during reproduction between an individual of genotype *j* and an individual of genotype *k*, depending on the recombination rate *ρ* between the color pattern locus *P* and the preference locus *M* (see Supp. S1 for detailed expression of *coef* (*i, j, k, ρ*)). We checked that for all *n* in {1, 2} the sum of *F*_*i,n*_ over all *i* is always equal to one, as expected (see Supp. S2).

### Model exploration

The complexity of this two-locus diploid model prevents comprehensive exploration with analytical methods, we therefore used numerical simulations to identify the conditions promoting the evolution of disassortative mating. All parameters and parameter intervals used in the different simulations are summarized in Table 2. The values of the basic predation mortality factor *d*_*m*_ and *d*_*n*−*m*_, the unpalatability *λ* and migration rate *mig* are chosen as conditions maintaining balanced polymorphism at the color pattern locus *P* assuming random mating, taken from (Joron and Iwasa, 2005).

Simulations are performed using Python v.3. and by using discrete time steps as an approximation (Euler method) (see Supp. S3 for more details about the numeric resolution). We checked that reducing the magnitude of the time step provided similar dynamics (see fig. S8), ensuring that our discrete-time simulations provide relevant outcomes. Note that all scripts used in this study are available on GitHub: https://github.com/Ludovic-Maisonneuve/Evolution_and_genetic_architecture_of_disassortative_mating.

#### Introduction of preference alleles

We assume that random mating is the ancestral preference behavior. Before introducing preference alleles, we therefore introduce color pattern alleles in equal proportions, and let the population evolves under random mating until the dynamical system reaches an equilibrium. We assume that a steady point is reached when the variation of genotype frequencies in the numerical solution during one time unit is below 10^−5^ (see Supp. S4 for more details). At this steady state, we then introduce the preference allele *dis* in proportion 0.01 (when exploring hyp. 1) or the preference alleles *m*_*a*_, *m*_*b*_, *m*_*c*_ in proportion 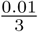 (when exploring hyp. 2).

After the introduction of preference alleles, we follow the evolution of disassortative mating and its consequences in the two populations:

- Early dynamic : First, we identify the range of parameters enabling the emergence of disassortative mating, by tracking genotype numbers during the first 100 time steps after the introduction of preference alleles.
- Steady state : Then, we study the long-term evolutionary outcome associated with the changes in mating behavior, by computing genotype numbers at equilibrium, *i.e*. by running simulations until the variation of genotype frequency during one time unit is below 10^−5^ (see Supp. 4 for more details).

#### Summary statistics

To facilitate the interpretation of our results, we compute a number of summary statistics from the outcomes of our simulations. We define haplotypes as the pairs of alleles in 𝒜_*P*_ × 𝒜_*M*_ containing two alleles located on the same chromosome or inherited from the same parent. We then calculate haplotype frequencies in patch 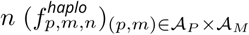 for *n* ∈ {1, 2}. Then for (*p, m, n*) ∈ 𝒜_*P*_ × 𝒜_*M*_ × {1, 2}, the frequency of haplotype (*p, m*) in patch *n* is given by:

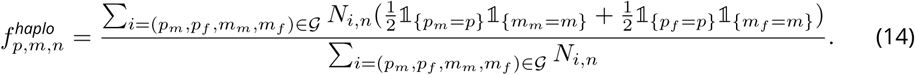

The estimation of haplotype frequencies allows to characterize the association between color pattern alleles and preference alleles, leading to different mating behaviors among partners with different color patterns, specifically under the *recognition/trait* hypothesis (Hyp.2). To characterize female mating preferences generated by the different genotypes at locus *M* and the link with their own color pattern phenotype, we then distinguish two main behaviors emerging under hyp. 2 (fig. S6 and S7) for *attraction* (hyp. 2.a) and *rejection* (hyp. 2.b) hypotheses respectively:

- Self-acceptance : females mate with males displaying their own color pattern phenotype.
- Self-avoidance : females do not mate with males displaying their own color pattern phenotype.

In order to compare the mating behaviors observed under *self-referencing* (hyp. 1) *attraction* (hyp. 2.a) and *rejection* (hyp. 2.b) hypotheses, we compute population statistics, *P*_*s-acc*_ (see equation (15)) and *P*_*s-av*_ (see equation (16)) as the proportion of individuals exhibiting respectively a self-acceptance or a self-avoidance behavior throughout both patches. These two inferred behaviors can be directly compared with mate preferences empirically estimated. For example, in experiments where females can choose partners among males displaying different color patterns (Chouteau, Llaurens, et al., 2017), the proportion of females mating with males displaying their own phenotype color pattern can be easily scored and compared to the proportion of self-accepting individuals computed in our model.

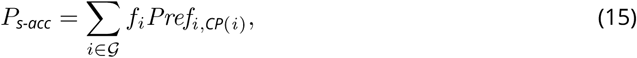

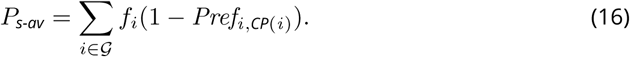

## Results

### Effect of mate choice on polymorphism

The emergence of disassortative mating requires initial polymorphism at the trait used as mating cue. Because the costs associated with mate searching and courting penalize females preferring rare phenotypes, the distribution of color pattern variation in the population may be an important condition for the emergence of disassortative mating. In turn, the evolution of disassortative mating is likely to generate a positive selection on rare phenotypes, therefore enhancing polymorphism at the color pattern locus *P*. To disentangle the feedbacks between polymorphism of the cue and evolution of disassortative mating, we first investigate the impact of different mating behaviors on the distribution of color pattern phenotypes within populations.

Under random mating, the frequencies of color pattern alleles at equilibrium computed for different migration rates *mig* show that polymorphism can be maintained through an equilibrium between spatially heterogeneous selection and migration (fig.1 (a)), consistent with previous results from the literature (Joron and Iwasa, 2005). In the absence of migration however, phenotypes *A* and *B* are fixed in the populations living in patch 1 and 2 respectively, owing to their mimetic advantage within their respective communities. Polymorphism with persistence of phenotypes *A* and *B* within each population can only be maintained with migration, but in all cases the non-mimetic phenotype *C* is not maintained in any of the two populations (fig.1 (a)).

To test for an effect of mate choice on this selection/migration equilibrium, we then compare those simulations assuming random mating (*i.e*. with preference alleles *r*) with simulations where *self-referencing* preference alleles generating either assortative (*sim* allele) or disassortative (*dis* allele) behavior were introduced at the mate choice locus *M* (hyp. 1), assumed to be fully linked to the color pattern locus *P* (*ρ* = 0). Assuming assortative mating via *self-referencing* (hyp. 1) the results are similar to whose observed under random mating (fig.1 (a),(b)). Nevertheless, the proportion of locally adapted alleles is higher than under random mating because assortative mating reinforces positive frequency dependent selection on those alleles. In contrast, disassortative mating maintains a higher degree of polymorphism, with the two mimetic phenotypes *A* and *B* and the non-mimetic phenotype *C* persisting within both populations, for all migration rates (fig.1 (c)). The non-mimetic phenotype *C* is rarely expressed because allele *c* is recessive. Nevertheless, individuals displaying phenotype *C* benefit from a high reproductive success caused by disassortative mating. Indeed, the strict disassortative preference assumed here strongly increases the reproductive success of individuals displaying a rare phenotype such as *C*. Negative frequency-dependent selection (FDS hereafter) on color pattern thus generated by disassortative mating counteracts the positive FDS due to predator behavior acting on the same trait. Therefore, disassortative mate preferences can strongly promote polymorphism within the two populations living in patch 1 and 2 respectively. When polymorphism is high, the cost of finding a dissimilar mate may be reduced, therefore limiting selection against disassortative preferences. Our results thus highlight the decreased cost of finding a dissimilar mate once disassortative mating becomes established.

### Linked genetic load favors the persistence of maladaptive alleles

In the following simulations, the migration parameter *mig* is set to 0.1, to allow for the persistence of polymorphism of color pattern phenotype *A* and *B* when assuming random mating. We then investigate the influence of a genetic load associated with the different color pattern alleles on polymorphism at the color pattern locus *P*, under random mating. This allows quantifying the effect of heterozygote advantage, independently of the evolution of mating preferences. We observe that the non-mimetic phenotype *C* is maintained together with phenotypes *A* and *B* within both populations, when (i) all three alleles carry a genetic load of similar strength, *i.e. δ*_*a*_ = *δ*_*b*_ = *δ*_*c*_ *>* 0 or (ii) when allele *c* is the only one without any associated genetic load (*δ*_*a*_ = *δ*_*b*_ *>* 0 and *δ*_*c*_ = 0) (fig. S9). In contrast, phenotype *C* is not maintained when a genetic load is associated with the non mimetic allele *c* only (*δ*_*a*_ = *δ*_*b*_ = 0 and *δ*_*c*_ *>* 0), or when this load is stronger than the one associated with alleles *a* and *b* (fig. S9). The heterozygote advantage generated by genetic load associated with the dominant mimetic alleles at locus *P* therefore favors the persistence of a balanced polymorphism and more specifically promotes the maintenance of allele *c* in both patches, even though this allele does not bring any benefit through local (mimicry) adaptation.

### Evolution of disassortative mating

Because we expect heterozygote advantage at the color pattern locus *P* to enhance the evolution of disassortative mating preferences at locus *M*, we first investigate the influence of a genetic load on the evolution of disassortative behavior by testing the invasion of *self-referencing* mutation triggering self-avoidance *dis* (hyp. 1) in a population initially performing random mating with genotype frequencies at equilibrium. We compute the frequency of mutants 100 time units after their introduction, assuming full linkage between loci *P* and *M*. Figure 2 shows that the genetic load associated with alleles *a* and *b* (*δ*_*a*_ = *δ*_*b*_), has a strong positive impact on the emergence of disassortative mating. The genetic load associated with the recessive allele *c* (*δ*_*c*_) has a weaker positive effect on the evolution of disassortative mating. Simulations assuming different relative cost of choosiness (*c*_*r*_) show a similar effect of associated genetic loads (see fig. 2). However the cost of choosiness reduces the range of genetic load values allowing the emergence of disassortative preference. When this cost is high, the invasion of mutant allele *dis* is prevented, regardless of the strength of genetic load (see fig. 2(d)). Although an increased cost of choosiness slows down the invasion of the disassortative mating mutant *dis* (see fig. 2), a genetic load linked to the color pattern locus *P* generally favors the emergence of disassortative mating in both patches.

**Figure 1.**
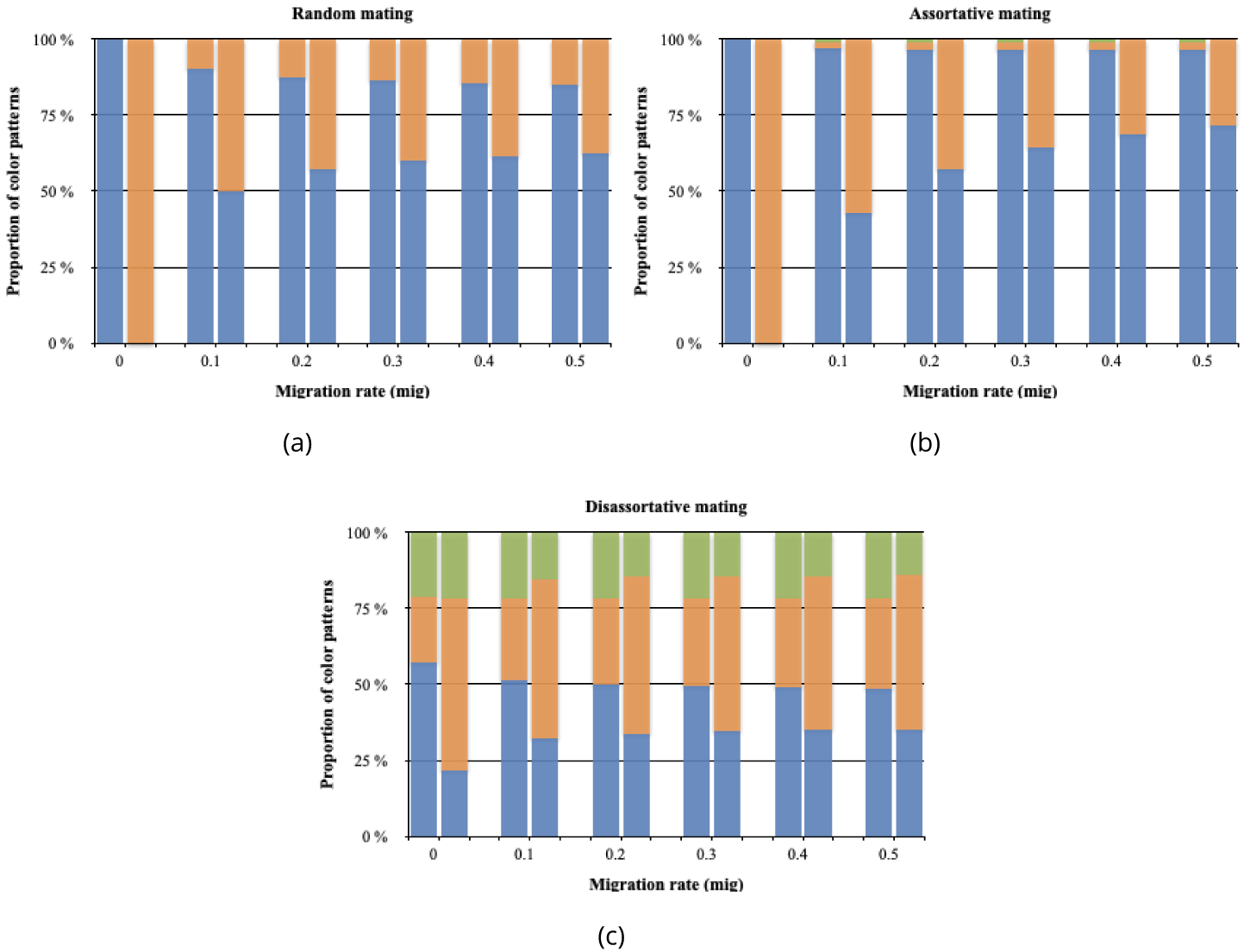
Influence of mate preferences on color pattern diversity within both patches. The equilibrium frequencies of color pattern phenotypes in patches 1 and 2 for different migration rates *mig* are computed assuming different mating behaviors, *i.e*., random (a), assortative (b) or disassortative (c). The heights of the colored stacked bars indicate the frequencies of color pattern phenotypes *A, B* and *C* (blue, orange and green areas respectively) in patches 1 and 2 (on the left and right side respectively, for each migration level). The three alleles at the locus *P* controlling color pattern variations are introduced in proportion 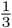 in each patch. The locus *M* controls for the *self-referencing* based mate preferences (hyp. 1): preferences alleles *r, sim* and *dis* were introduced in simulations shown in panel (a), (b) and (c) respectively. Simulations are run assuming *r* = 1, *K* = 2000, 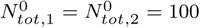,*λ* = 0.0002, *d*_*m*_ = 0.05, *d*_*n−m*_ = 0.15, *ρ* = 0, *c*_*r*_ = 0.1, *δ*_*a*_ = *δ*_*b*_ = *δ*_*c*_ = 0 and *δ* = 0.1.

**Figure 2.**
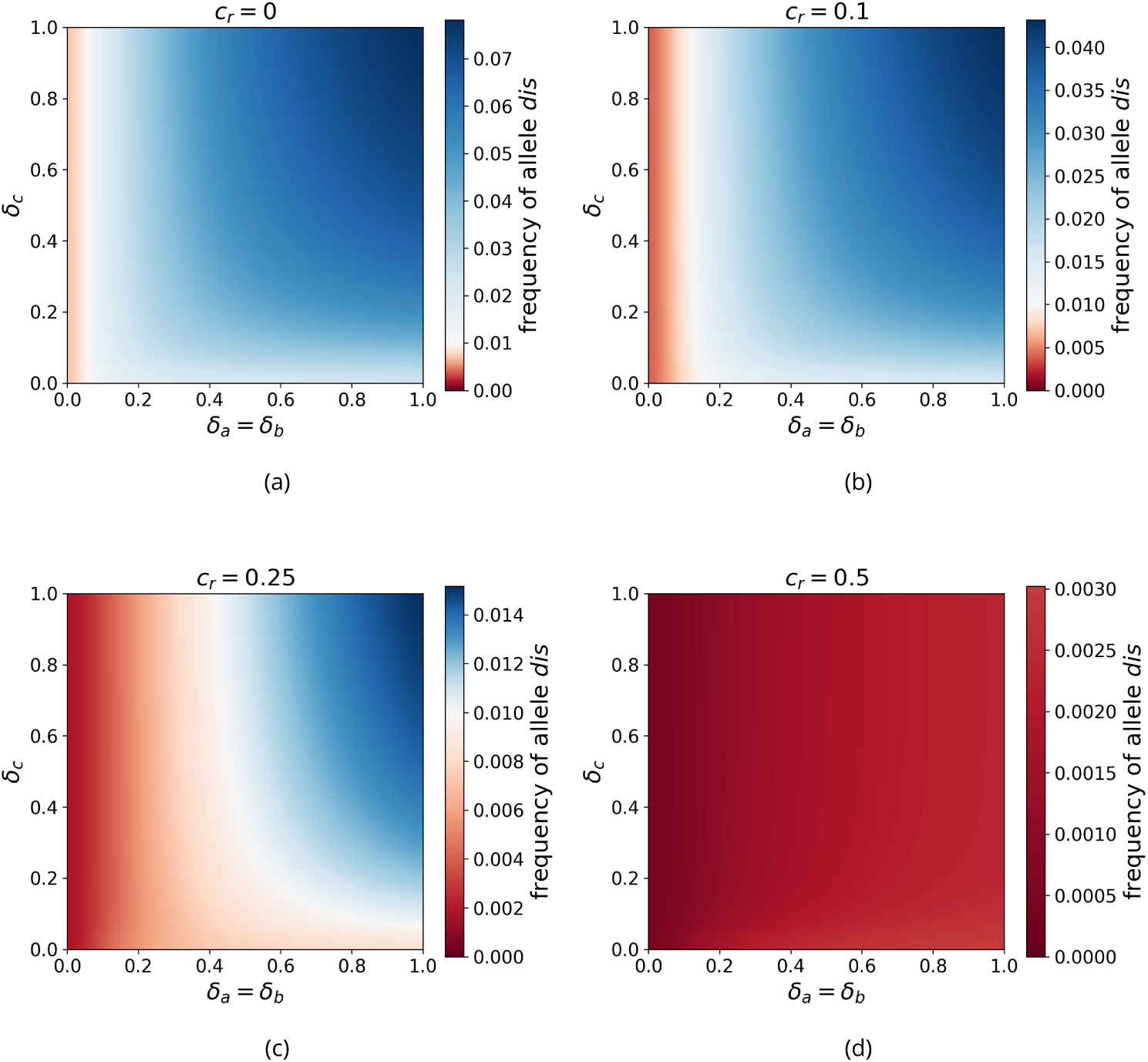
Influence of a linked genetic load on the emergence of disassortative mating for different costs of choosiness, assuming *self-referencing* (hyp. 1). The frequency of the mutant allele *dis* is shown 100 time units after its introduction depending on the strength of genetic load associated with the dominant alleles *a* and *b* (*δ*_*a*_ = *δ*_*b*_) and to the recessive allele *c, δ*_*c*_. The initial frequency of allele *dis* was 0.01, the area where mutant allele increase (resp. decrease) is shown in blue (resp. red). Simulations are run assuming either (a) no cost of choosiness *c*_*r*_ = 0, (b) a low cost of choosiness *c*_*r*_ = 0.1, (c) an intermediate cost of choosiness *c*_*r*_ = 0.25 or (d) an elevated cost of choosiness *c*_*r*_ = 0.5. Simulations are run assuming *r* = 1, *K* = 2000, 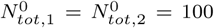, *λ* = 0.0002, *d*_*m*_ = 0.05, *d*_*n−m*_ = 0.15, *mig* = 0.1 and *ρ* = 0.

To investigate the long-term evolution of disassortative mating promoted by the genetic loads associated with color pattern alleles, we then compute the frequency of mutant allele *dis* at equilibrium in conditions previously shown to promote its emergence (*i.e*. assuming limited cost of choosiness). Figure 3 shows that the mutant preference allele *dis* is never fixed within populations. This suggests that the heterozygote advantage at locus *P* allowing the emergence of disassortative mating decreases when this behavior is common in the population. The *dis* mutant nevertheless reaches high frequencies when the genetic load associated with the recessive allele *c* is intermediate (*δ*_*c*_ ≈ 0.35) and the genetic load associated with dominant alleles *a* and *b* is strong (see fig. 3). This result seems surprising because the highest level of disassortative mating is not reached when the genetic load is at the highest in all the three alleles at locus *P*. On the contrary, disassortative mating is favoured when a genetic load is associated with the dominant alleles only: disassortative mating limits more the cost of producing unfit offspring when a genetic load is associated with dominant alleles, because these alleles are always expressed as color pattern phenotypes, and therefore avoided by females with disassortative preferences.

**Figure 3.**
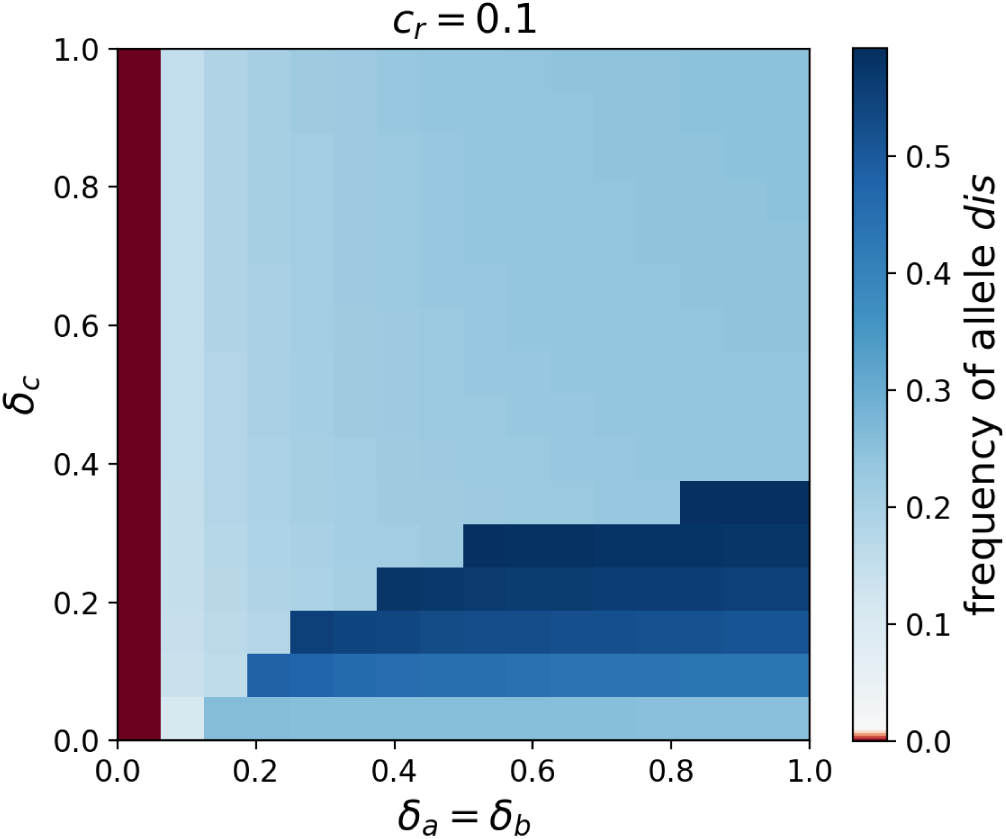
Influence of a linked genetic load on the level of disassortative mating at equilibrium for low cost of choosiness (*c*_*r*_ = 0.1), assuming *self-referencing* (hyp. 1). The frequency of the mutant allele *dis* is shown at equilibrium after its introduction depending on the strength of genetic load associated with the dominant alleles *a* and *b* (*δ*_*a*_ = *δ*_*b*_) and with the recessive allele *c, δ*_*c*_. The initial frequency of allele *dis* is 0.01. The area where the frequency of the mutant allele increases (resp. decrease) is shown in blue (resp. red). Simulations are run assuming *r* = 1, *K* = 2000, 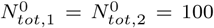, *λ* = 0.0002, *d*_*m*_ = 0.05, *d*_*n−m*_ = 0.15, *mig* = 0.1, *ρ* = 0 and *c*_*r*_ = 0.1.

### How does the genetic architecture of mating preference influence the evolution of disassortative mating ?

To study the impact of the genetic architecture of mate preferences on the evolution of disas-sortative mating, we then compare the invasion of *self-referencing* alleles *dis* with the invasion of *recognition/trait* alleles (*i.e*. alleles *m*_*r*_, *m*_*a*_, *m*_*b*_ and *m*_*c*_ controlling random mating and specific recognition of phenotype *A, B* and *C* respectively, hyp. 2). We assume loci *P* and *M* to be fully linked (*ρ* = 0), and compare simulations where mate preference alleles trigger either disassortative preference (hyp. 1), *attraction* (hyp. 2.a) or *rejection* (hyp. 2.b) of the recognized color pattern phenotype. We report the frequencies of haplotypes, in order to follow the association of color pattern and preference alleles (fig.4(a), fig.4(b) and fig.4(c) respectively).

Under a *self-referencing* rule, alleles *a* and *b* are associated with preference allele *dis* as soon as the genetic load associated with the dominant alleles (alleles *a* and *b*) is greater than 0. Indeed disassortative mating favors the production of heterozygotes and reduces the expression of the genetic load in the offspring. In contrast, the non-mimetic allele *c*, not associated with any genetic load, is preferentially linked with the random mating allele *r*. This result is surprising because heterozygotes carrying a *c* allele have a lower predation risk than homozygotes with two *c* alleles: homozygotes are indeed non-mimetic in both patches, while heterozygotes are mimetic in one out of the two patches. However, the benefit associated with haplotype (*c, dis*) through increased production of heterozygous offspring is weak. Because of the genetic load associated with the dominant color alleles *a* and *b, c* allele is common in the population, resulting in relatively high frequency of homozygotes with two *c* alleles, and of heterozygotes with one *c* allele. Alleles *a* and *b* are frequently linked with the disassortative preference allele *dis*, further promoting the formation of heterozygotes. Since *c* allele is recessive, disassortative crosses between individuals with phenotype *C* and either *A* or *B* then frequently produce progeny with half of the offspring carrying two *c* alleles, suffering from increased predation. The limited survival of these offspring reduces the benefits associated with the haplotype (*c, dis*). Because the *dis* allele is also associated with a cost of choosiness, linkage between allele *c* and the random mating allele *r* could then be promoted.

When preference alleles cause female attraction to males exhibiting a given phenotype (hyp. 2.a), only haplotypes (*a, m*_*c*_) and (*c, m*_*a*_) are maintained in both patches at equilibrium (fig.4(b)). The haplotype (*a, m*_*c*_) benefits from both positive selection associated with mimicry and limited expression of the genetic load due to the preferential formation of heterozygotes. Haplotype (*c, m*_*a*_) is maintained because of the benefit associated with the choice of the most frequent mimetic phenotype *A*, and the limited expression of the non-mimetic phenotype *C* due to *c* being recessive. The proportion of haplotype (*a, m*_*c*_) decreases as the genetic load associated with allele *a* increases. Indeed the mating between two individuals of genotype (*a, c, m*_*c*_, *m*_*a*_) becomes more likely and leads to the formation of individuals (*a, a, m*_*c*_, *m*_*c*_) suffering from the expression of the genetic load. Allele *b* is then lost because of the dominance relationships between alleles *a* and *b*. Phenotype *A* is more commonly expressed than phenotype *B*: haplotype (*c, m*_*a*_) is thus favoured over haplotype (*c, m*_*b*_), through increased mate availability. Sexual selection caused by disassortative preferences generate a strong disadvantage associate with *b* allele, ultimately leading to its extinction.

By contrast, when mate preference is based on alleles causing *rejection* behavior (hyp. 2.b) and when a genetic load is associated with the mimetic alleles *a* and *b* at locus *P*, these alleles become associated with the corresponding rejection alleles at locus *M* (*i.e*. (*a, m*_*a*_) and (*b, m*_*b*_) have an intermediate frequencies in both patches) (fig.4(c)). Non mimetic allele *c* becomes associated with random mating preference allele *r*. The three alleles (*a, b* and *c*) persist within patches for all positive values of genetic load. This contrasts with the evolutionary outcome observed under attraction rule (hyp. 2.a) where mimetic allele *b* is lost if the genetic load is greater than 0 (fig. 4(b)).

**Figure 4.**
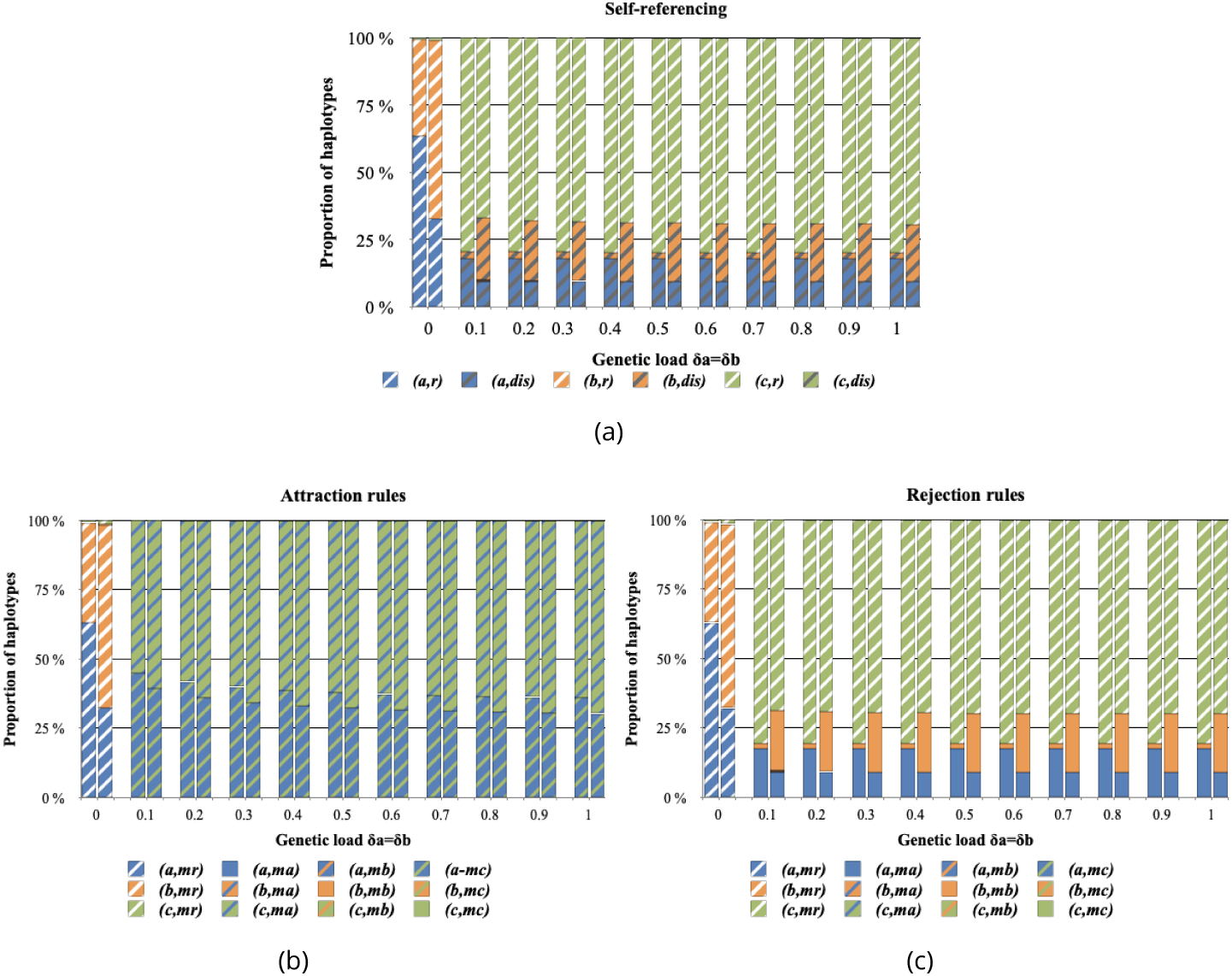
Influence of a genetic load on haplotype diversity, assuming (a) *self-referencing* (hyp. 1), (b) *attraction* rule (hyp. 2.a) or (c) *rejection* rule (hyp. 2.b) at the preference locus (*recognition/trait*). The proportion of haplotypes at equilibrium after the introduction of preference alleles in both patches are shown for different values of genetic load associated with alleles *a* and *b* (*δ*_*a*_ = *δ*_*b*_). For each value of genetic load (*δ*_*a*_ = *δ*_*b*_) the first and second bars show the frequencies of haplotypes in the patches 1 and 2 respectively. Simulations are run assuming *r* = 1, *K* = 2000, 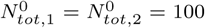, *λ* = 0.0002, *d*_*m*_ = 0.05, *d*_*n−m*_ = 0.15, *ρ* = 0, *mig* = 0.1, *δ*_*c*_ = 0, *δ* = 0.1 and *c*_*r*_ = 0.1.

We then investigate how these haplotype frequencies translate into individual behaviors in the populations at equilibrium. As highlighted in fig.5, the proportion of each behavior depends more on the existence of a genetic load linked to dominant alleles, than on its strength. The proportion of disassortative mating is similar when assuming *self-referencing* (hyp. 1) and *recognition/trait* leading to rejection (hyp. 2.b) (*P*_*s-av*_ ≈ 48%) (fig.5(a) and 5(c)).

By contrast, when we consider preference alleles leading to *attraction* (hyp. 2.a), the disassortative behavior is scarcer at equilibrium (*P*_*s-av*_ ≈ 36%) (fig. 5(b)). This may seem surprising given that most haplotypes are formed by a color pattern allele linked with an *attraction* allele for a different color pattern (fig. 4(b)). Nevertheless, the color pattern allele *c* is linked to *m*_*a*_ coding for attraction to *A*. As a consequence, most individuals formed are heterozygous at both the color pattern locus *P* (with one allele *a* and one allele *c*) and at the preference locus *M* (with one preference allele coding for attraction toward phenotype *A* and another preference allele triggering attraction toward phenotype *C*). These double heterozygotes thus benefit from mimicry and avoid the expression of deleterious mutations, and are self-accepting. However, under the *self-referencing* (hyp. 1) or *rejection* (hyp. 2.b) rules disassortative mating is more likely to emerge. Indeed under hyp. 2.b, haplotypes composed by a phenotype allele and its corresponding preference allele ((*a, m*_*a*_) for example) generally immediately translates into a self-avoiding behavior, whatever the genotypic combinations within individuals. Moreover under hyp. 1 disassortative haplotype, *i.e*. an haplotype where the preference allele is *dis*, always generates a disassortative behavior.

**Figure 5.**
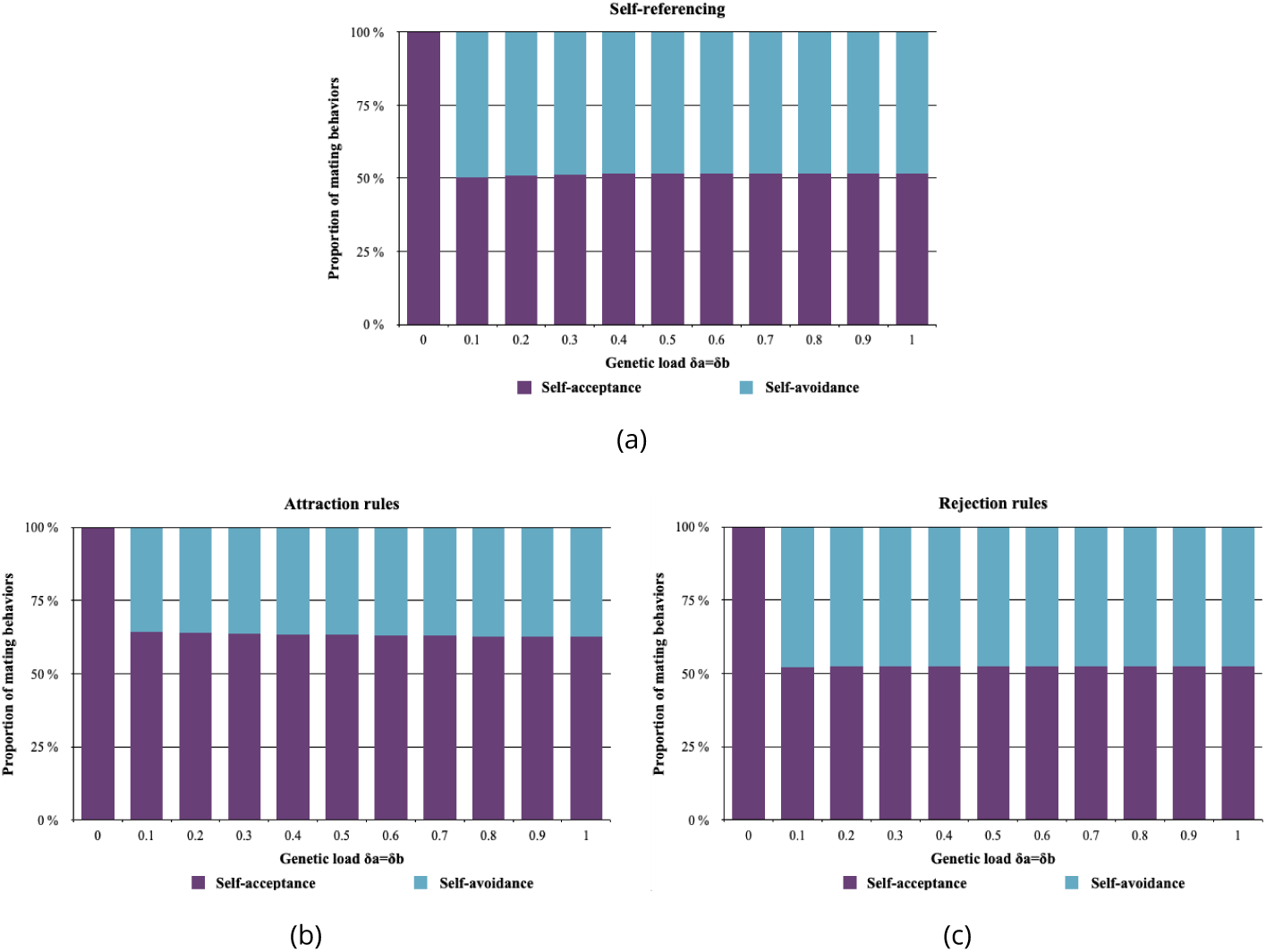
Influence of a genetic load on the distribution of mating behavior observed at the population level, assuming (a) *self-referencing* (hyp. 1), (b) *attraction* rule (hyp. 2.a) or (c) *rejection* rule (hyp. 2.b) at the preference locus (*recognition/trait*). The proportion of individuals displaying self-acceptance *P*_*s-acc*_ (in purple) and self-avoidance *P*_*s-av*_ (in blue) obtained at equilibrium after the introduction of preference alleles are shown for different values of the level of genetic load of *δ*_*a*_ and *δ*_*b*_. Simulations are run assuming *r* = 1, *K* = 2000, 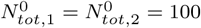, *λ* = 0.0002, *d*_*m*_ = 0.05, *d*_*n−m*_ = 0.15, *ρ* = 0, *mig* = 0.1, *δ*_*c*_ = 0, *δ* = 0.1 and *c*_*r*_ = 0.1.

This highlights that the genetic architecture of mate preference plays a key role in the evolution of the mating behavior of diploid individuals: the evolution of disassortative haplotypes inducing disassortative preferences do not necessarily cause disassortative mating at the population level. At equilibrium, the proportion of self-avoidance behavior in the population hardly depends of the strength of the genetic load (figure 5). However, the strength of the genetic load does increase the speed of evolution of disassortative mating (see fig. S10 comparing the invasion dynamics of the self-avoiding behavior when assuming different levels of genetic load), therefore suggesting stronger positive selection on disassortative mating when the genetic load associated with dominant wing color pattern alleles is higher.

### Impact of linkage between loci *P* and *M* on the evolution of disassortative mating

In previous sections, we observed that the genetic load associated with the two most dominant alleles at the color pattern locus *P* impacts the evolution of mate choice. Assuming that the color pattern locus *P* and the preference locus *M* are fully linked, we also noticed that disassortative mating is more prevalent at equilibrium under the *self-referencing* rule (hyp. 1) and the *rejection rule* (hyp. 2.b) rather than under the *attraction* (hyp. 2.a) rule. We then test for an effect of recombination between alleles at the two loci on the evolution of mate choice by performing simulations with different values of the recombination rate *ρ*.

Assuming *self-referencing* (hyp. 1), increasing recombination rate strongly promotes the self-avoidance behavior (*P*_*s-av*_ ≈ 98%) (see fig. 6(a)). Selection generated by the genetic load associated to color pattern alleles *a* and *b* promotes their linkage with the disassortative *self-referencing* allele *dis*, while the genetic-load free allele *c* tends to be linked to the random mating allele *r* (as observed in simulations assuming no recombination, fig. S11(a)). Because allele *dis* reaches a high frequency in the population, recombination generates a large density of recombinant haplotypes (*a, r*), (*b, r*), (*c, dis*). Haplotypes (*a, r*) and (*b, r*) are disfavored because they lead to a the production of offspring suffering from the expression of a genetic load, whereas (*c, dis*) leads to the production of viable offspring. Therefore, under the *self-referencing* hypothesis (hyp. 1), recombination thus significantly increases the proportion of disassortative mating.

**Figure 6.**
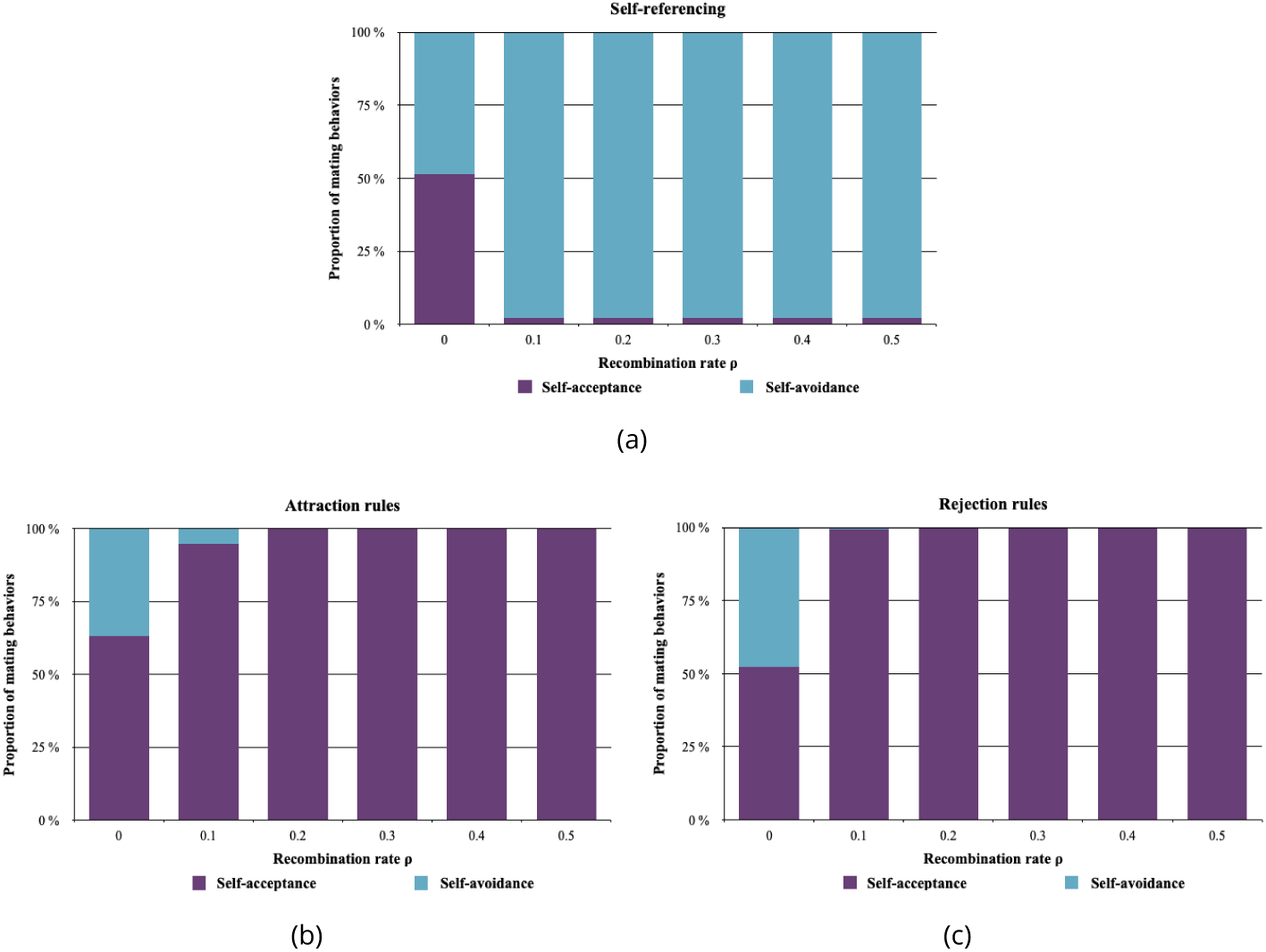
Influence of the recombination rate between color pattern and preference alleles on the distribution of mating behavior observed at the population level, assuming different genetic architectures of mate preferences: either (a) *self-referencing* (hyp. 1), or *recognition/trait* leading to (b) *attraction* rule (hyp. 2.a) or (c) *rejection* rule (hyp. 2.b). The proportion of individuals displaying self-acceptance *P*_*s-acc*_ (in purple) and self-avoidance *P*_*s-av*_ (in blue) obtained at equilibrium are shown for different values of recombination rate *ρ* between the preference locus *M* and the color pattern locus *P*. Simulations are run assuming *r* = 1, *K* = 2000, 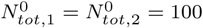, *λ* = 0.0002, *d*_*m*_ = 0.05, *d*_*n−m*_ = 0.15, *mig* = 0.1, *δ*_*a*_ = *δ*_*b*_ = 0.5, *δ*_*c*_ = 0, *δ* = 0.1 and *c*_*r*_ = 0.1.

Under *self-referencing* rule (hyp. 1), mate preference depends on the phenotype displayed by the individual, so that allele *dis* always translates into a disassortative behavior. By contrast, when assuming *recognition/trait* for a given color pattern allele (hyp. 2), mating behavior depends only on the genotype at the preference locus *M*, independently from the color pattern of the female. We therefore expect a stronger effect of recombination rate on mate choice evolution. Figure 6 indeed confirms this prediction. Under *attraction* (hyp. 2.a) and *rejection* (hyp. 2.a) rules, the most striking effect is observed when comparing simulations assuming *ρ* = 0 *vs ρ >* 0: self-avoidance behavior is scarcely observed in the population (*P*_*s-av*_ ≈ 1%) when there is recombination (*ρ >* 0).

Our results suggest that disassortative mating can emerge either (1) under the *self-referencing* rule or (2) under the *recognition/trait* rule assuming a tight linkage between the loci controlling cue and preference. Nevertheless, strict *self-referencing* behaviour, under which preference varies according to the chooser’s phenotype, is scarcely observed in natural populations (see Kopp et al., 2018 for a review). We thus expect that disassortative mating might emerge when the mating cue and the preference loci are tightly linked or are controlled by a single pleiotropic gene.

## Discussion

### Genetic architecture of disassortative mating: theoretical predictions

Our model shows that without recombination between color pattern (locus *P*) and preference alleles (locus *M*), disassortative mating is more likely to emerge when the genetic architecture is with *self-referencing* (hyp. 1) or with color pattern recognition triggering *rejection* (hyp. 2.b). When preference alleles cause *attraction* to males exhibiting a given phenotype (hyp. 2.a), heterozygote advantage favors haplotypes formed by a color pattern allele linked with an attraction allele for a different color pattern. However, these haplotypes do not necessarily imply a complete self-avoidance behavior in females carrying them. The co-dominance assumed at the preference locus indeed generates preference for two different phenotypes in heterozygotes at the locus *M*, favoring self-acceptance. This effect is reinforced by the mate choice, promoting the association between a color allele and the corresponding attraction allele in the offspring, and therefore increasing the emergence of self-accepting genotypes. This might explain the low proportion of self-avoidance behavior observed within populations, when assuming the *attraction* rule (hyp. 2.a). By contrast, when recombination between the two loci does occur, a *self-referencing* architecture (hyp. 1) may facilitate the evolution of disassortative mating. The genetic basis of disassortative mating is largely unknown in natural populations. Assortative mating is better documented, for instance in *Heliconius* butterflies where it is generally associated with attraction towards a specific cue. The locus controlling preference for yellow *vs*. white in *H. cydno* maps close to the gene *aristaless*, whose expression differences determine the white/yellow switch in this species (Kronforst et al., 2006; Westerman et al., 2018). In *H. melpomene*, a major QTL associated with preference towards red was identified in crosses between individuals displaying a red pattern and individuals with a white pattern (Merrill et al., 2019). This QTL is also located close to the gene *optix* involved in the variation of red patterning in *H. melpomene*. Assortative mating in *Heliconius* thus seems to rely on alleles encoding preference for specific cues, linked to with loci involved in the variation of these cues. Similarly, our model suggests that the genetic architecture of disassortative mating might involve tight linkage between the cue and the preference loci. However, in contrast with the attraction alleles documented in species where assortative mating behavior is observed, our results show that alleles coding for rejection toward certain cue are more likely to promote the evolution of disassortative mating.

Similar mate preference is obtained with some *recognition/trait* (hyp.2) genotypes than with some *self-referencing* (hyp. 1) genotypes: for example, under the *rejection* rule (hyp. 2.b), the genotype (*a, a, m*_*a*_, *m*_*a*_) leads to the same mate preference as the genotype (*a, a, dis, dis*) under the *self-referencing* genetic architecture. Introducing recombination in the *recognition/trait* architecture then enables the decoupling of the mating cue and of its corresponding preference alleles, thereby disrupting the self rejection behavior. Furthermore, under the *recognition/trait* architecture, our model distinguishes whether the specific recognition of the cue leads to *rejection* or *attraction*, and highlights that these two hypotheses lead to the evolution of different preference regimes: disassortative mating is more likely to emerge assuming a *rejection rule*. This rule indeed generates a greater density of self-rejecting haplotypes than the *attraction* rule, although recombination limits this effect.

The dominance relationships assumed at both the cue and preference loci are likely to impact our predictions on the evolution of disassortative mating. Disassortative mating is at an advantage when it favors the production of offspring that do not express the genetic load. Dominance relationships at the color pattern locus *P* signal the genetic load associated with dominant cue alleles. This explains why disassortative mating is favored when the genetic load is low in the recessive cue alleles and large in dominant cue alleles. The co-dominance assumed at the preference locus generates preferences toward two different phenotypes in heterozygotes at the preference locus. We suspect that alternative hypotheses on dominance at the preference locus may modulate our predictions on the evolution of disassortative mating.

Altogether, our theoretical model shows that the genetic basis of mate preferences has a strong impact on the evolution of disassortative mating at loci under heterozygote advantage. This emphasizes the need to characterize the genetic basis of mate preference empirically and the linkage disequilibrium with the locus controlling variation in the mating cues.

### Evolution of disassortative mating results from interactions between dominance and deleterious mutations

Here, we confirm that the evolution of disassortative mating is promoted by the heterozygote advantage associated with alleles determining the mating cue. As mentioned above, the phenotype of the chosen individuals depends on the dominance relationships at the color pattern locus. Our model highlights that a genetic load associated with the dominant alleles contributes more to disassortative mating than a genetic load associated with the most recessive haplotype. This theoretical prediction is in accordance with the few documented cases of polymorphism promoted by disassortative mating. In the polymorphic butterfly *Heliconius numata* for instance, the top dominant haplotype *bicoloratus* is associated with a strong genetic load (Jay et al., 2019). Similarly, in the white throated sparrow, the dominant *white* allele is also associated with a significant genetic load (Tuttle et al., 2016). Again, in the self-incompatibility locus of the *Brassicaceae*, dominant haplotypes carry a higher genetic load than recessive haplotypes (Llaurens, Gonthier, et al., 2009). Disassortative mating is beneficial because it increases the number of heterozygous offspring with higher fitness. Once disassortative mating is established within a population, recessive deleterious mutations associated with the dominant haplotype become sheltered because the formation of homozygotes carrying two dominant alleles is strongly reduced, thereby limiting the opportunities for purging via recombination (Llaurens, Gonthier, et al., 2009). Falk and Li, 1969 proved that disassortative mate choice promotes polymorphism, and therefore limits the loss of alleles under negative selection. Disassortative mating might thus shelter deleterious mutations linked to dominant alleles, and reinforce heterozygote advantage. The sheltering of deleterious mutations is favored by the interaction between two aspects of the genetic architecture: dominance at the mating cue locus and limited recombination. This is likely to happen in polymorphic traits involving chromosomal rearrangements, where recombination is limited. Many rearranged haplotypes are indeed associated with serious fitness reduction as homozygotes (Faria et al., 2019), such as in the derived haplotypes of the supergene controlling plumage and mate preferences in the white-throated sparrow (Thomas et al., 2008). The deleterious elements in the inverted segment can be due to an initial capture by the inversions (Kirkpatrick, 2010), but they could also accumulate through time, resulting in different series of deleterious mutations associated to inverted and non-inverted haplotypes (Berdan et al., 2019).

Here, we assume that mate choice relied purely on a single cue. Nevertheless, mate choice could be based on other cues, controlled by linked loci and enabling discrimination between homozygotes and heterozygotes, thereby further increasing the proportion of heterozygous offsprings with high fitness. We also modelled strict preferences regarding color patterns, but choosiness might be less stringent in the wild, and may limit the evolution of disassortative mating. Depending on the cues and dominance relationships among haplotypes, different mate choice behaviors may also evolve, which might modulate the evolution of polymorphism within populations. Our model thus stresses the need to document dominance relationships among haplotypes segregating at polymorphic loci, as well as mate choice behavior and cues, to understand the evolutionary forces involved in the emergence of disassortative mating.

## Conclusions

Inspired by a well-documented case of disassortative mating based on cues subject to natural selection, our model shows that heterozygote advantage is likely to favor the evolution of disassortative mating preferences. We highlight that disassortative mating is more likely to emerge when loci code for self-referencing disassortative preference or rejection of specific cues. However rejection locus only promotes disassortative mating when they are in tight linkage with the locus controlling mating cue variation.

## Supplementary material

Script and codes are available online: https://github.com/Ludovic-Maisonneuve/Evolution_and_genetic_architecture_of_disassortative_mating

## Acknowledgments

The authors would like to thank Charline Smadi and Emmanuelle Porcher for feedbacks on the modeling approach developed here. We also thank Thomas Aubier and Richard Merrill and the whole Heliconius group for stimulating discussion on mate choice evolution in our favorite butterflies. We are also grateful to Roger Butlin, Charles Mullon, Tom Van Dooren and two anonymous reviewers for their thoughtful review of a previous versions of this manuscript. We also thank Sylvain Gerber for his help in improving the English of this article. This work was supported by the Emergence program from Paris City Council to VL and the ANR grant SUPERGENE to VL and MJ. Version 9 of this preprint has been peer-reviewed and recommended by Peer Community In Evolutionary Biology (https://doi.org/10.24072/pci.evolbiol.100109).

## S1: Mendelian segregation

To compute the proportion of a given genotype in the progeny of the different crosses occurring in the population, we define a function *coef*(*g*^*O*^, *g*^*M*^, *g*^*F*^, *ρ*) summarizing the Mendelian segregation of alleles assuming two diploid loci and a rate of recombination *ρ* between these loci. Let 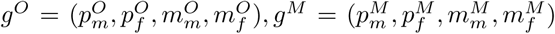 and 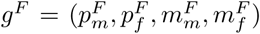 be the offspring, maternal and paternal genotypes respectively, all in 𝒢. For *I* ∈ {*O, M, F*},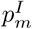 and 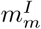 (resp. 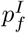 and 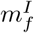) are the alleles on the maternal (resp. paternal) chromosomes. *coef*(*g*^*O*^, *g*^*M*^, *g*^*F*^, *ρ*) is the average proportion of genotype *g*^*O*^ in the progeny of a mother of genotype *g*^*M*^ mating with a father of genotype *g*^*F*^ given a recombination rate *ρ*.

Each diploid mother can produce four types of haploid gametes containing alleles 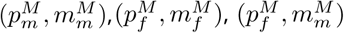or 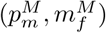, in proportion 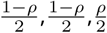 and 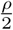 respectively. Then the proportion of gametes with alleles (*p, m*) ∈ 𝒜_*P*_ × 𝒜_*M*_ produced by the mother is given by the function *coef*_*haplotype*_(*p, m, g*^*M*^, *ρ*), where

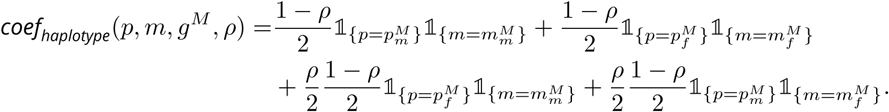

Similarly, each diploid father can produce four types of haploid gametes. The proportion of genotype (*p, m*) ∈ 𝒜_*P*_ × 𝒜_*M*_ in the gametes of a given father is given by the function *coef*_*haplotype*_(*p, m, g*^*F*^, *ρ*).

The average proportion of genotype *g*^*O*^ in the progeny of a cross between a mother of genotype *g*^*M*^ and a father of genotype *g*^*F*^ given a recombination rate *ρ* is given by:

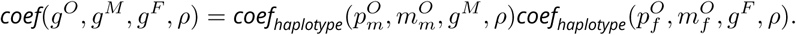

## S2: Checking of the computed genotype frequencies in the progeny of all crosses

We then check that the sum of the computed frequencies of the different genotypes *i* in the progeny of all crosses occuring in patch *n* ((*F*_*i,n*_))_*i*∈𝒢_ for *n* ∈ {1, 2}) actually equals to one. Let *n* be in {1, 2}, we have:

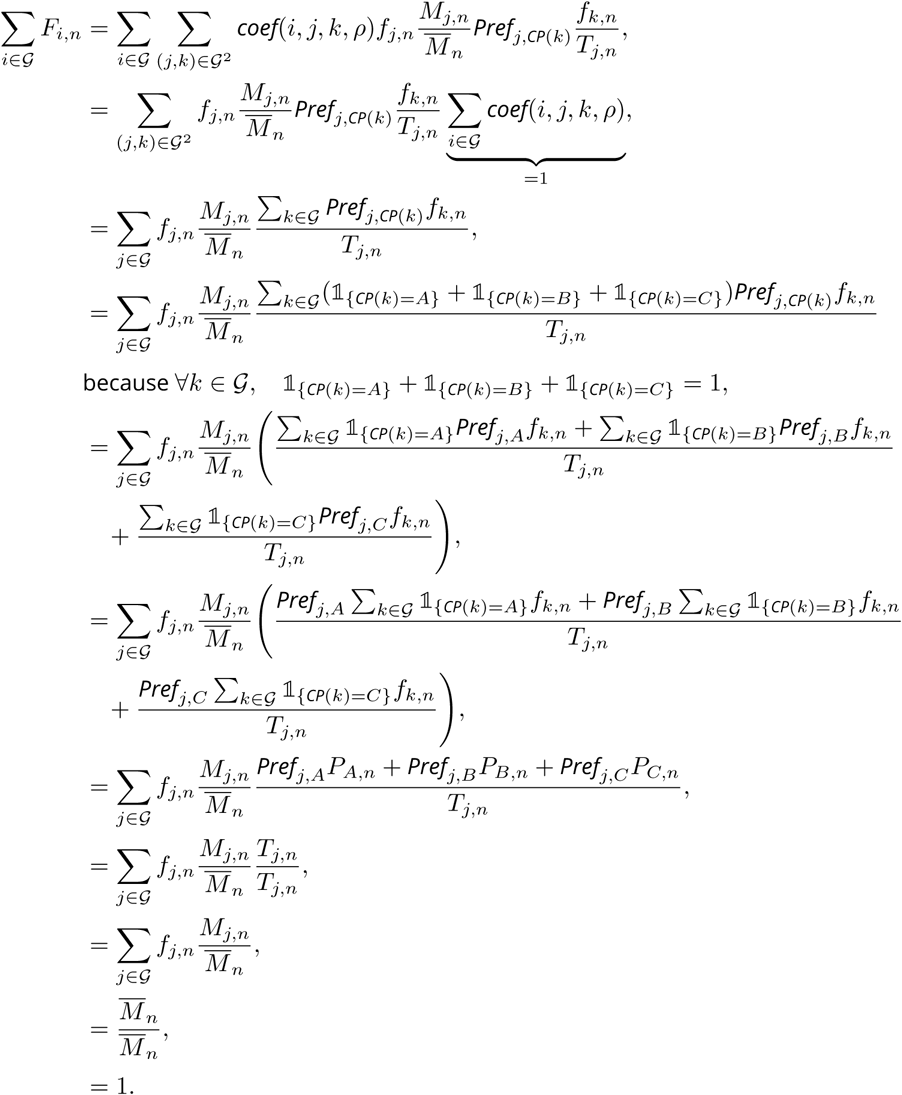

## S3: Numerical resolution

In this study, we used a numerical scheme to simulate our dynamical system. For (*i, n*) ∈ 𝒢 ×{1, 2}, let 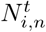 be the numerical approximation of *N*_*i,n*_(*t*). We use a explicit Euler scheme, therefore we approximate the quantity 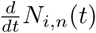 by

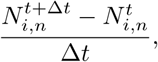

with Δ*t* being the step time in our simulations.

For (*i, n*) ∈ 𝒢 × {1, 2}, an approximation of equation 1 becomes:

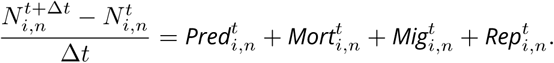

This equation is equivalent to:

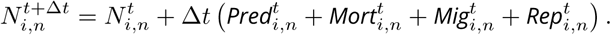

Given 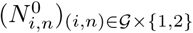, we can simulate an approximation of the dynamical system.

## S4: Numerical approximation of equilibrium states

To estimate the equilibrium reached by our dynamical system using simulations assuming different initial conditions, we define the variable *Var*^*t*^ quantifying the change in the numerical solution :

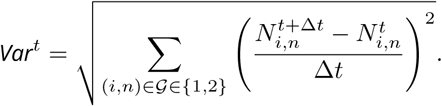

When 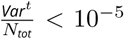, we assume that the dynamical system has reached equilibrium, with *N*_*tot*_ being the total density in both patches.

## Supplementary Figures

**Figure S5.**
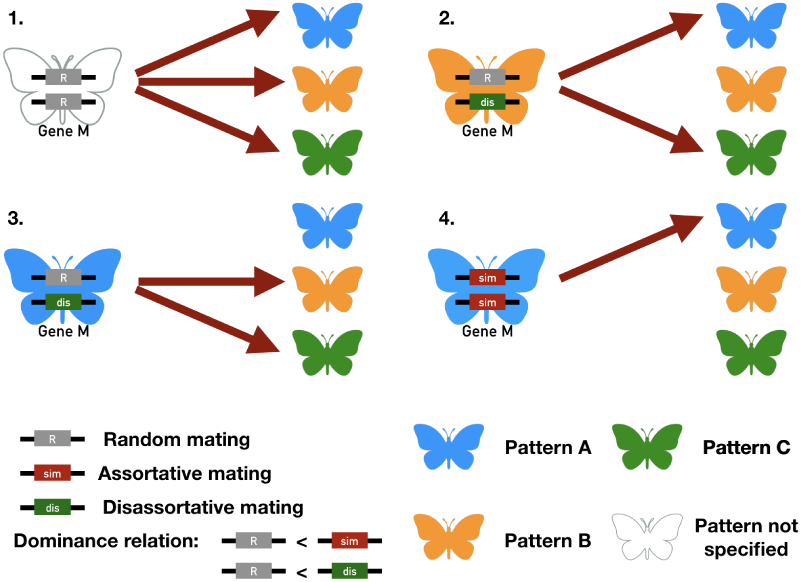
Mate preferences expressed by individuals carrying different genotypes at the preference locus *M*, assuming *self-referencing* (hyp. 1). 1. Butterflies carrying two *r* alleles mate at random, independently of either their own color pattern or the color pattern displayed by mating partners. 2-3. Butterflies carrying a *dis* allele display disassortative mating, and mate preferentially with individuals with a color pattern different from their own. 4. Butterflies carrying a *sim* allele display an assortative mating behavior and therefore preferentially mate with individuals displaying the same color pattern. Cases 1 and 4 therefore lead to *self-acceptance*, while cases 2 and 3 lead to *self-avoidance*.

**Figure S6.**
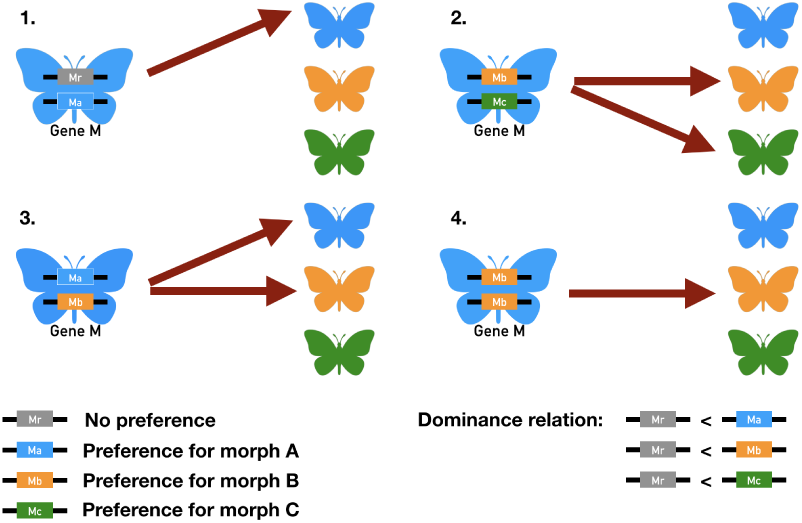
Mate preferences expressed by individuals carrying different genotypes at the preference locus *M*, assuming preference alleles encoding for *attraction* of specific color patterns (*recognition/trait*) (hyp. 2.a). 1. A butterfly displaying phenotype *A* (in blue) carries one allele coding for specific attraction toward partners displaying phenotype *A* (in blue) and the allele coding for random mating at the locus M controlling the mate choice. This butterfly will mate preferentially with individuals displaying phenotype *A*, resulting in assortative mating. 2. A butterfly displaying phenotype *A* (in blue) carries one allele coding for specific attraction toward partner displaying phenotype *B* (in orange) and one allele coding for specific attraction toward partners displaying phenotype *C* (in green). This individual will preferentially mate with individuals displaying phenotype *B* and *C*, resulting in disassortative mating 3. A butterfly displaying phenotype *A* (in blue) carries one allele coding for specific attraction toward partner displaying phenotype *A* (in blue) and one allele coding for specific attraction toward partners displaying phenotype *B* (in orange). This individual will preferentially mate with individuals displaying phenotype *A* and *B* 4. A butterfly displaying phenotype *A* (in blue) carries two alleles coding for specific attraction toward partner displaying phenotype *B* (in orange). This individual will preferentially mate with individuals displaying phenotype *B*, resulting in disassortative mating. Cases 1 and 3 therefore lead to *self-acceptance*, while cases 2 and 4 lead to *self-avoidance*.

**Figure S7.**
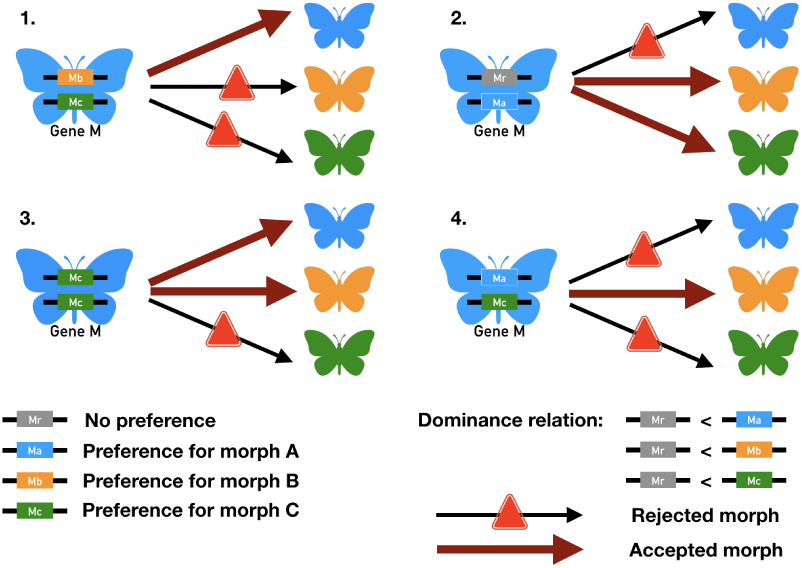
Mate preferences expressed by the different individuals carrying different genotypes at the preference locus *M*, assuming preference alleles encoding for *rejection* of specific color patterns (*recognition/trait*) (hyp. 2.a). 1. A butterfly displaying phenotype *A* (in blue) carries one allele coding for specific rejection toward partners displaying phenotype *B* (in orange) and one allele coding for specific rejection toward partners displaying phenotype *C* (in orange). This butterfly will mate preferentially with individuals displaying phenotype *A*, resulting in assortative mating. 2. A butterfly displaying phenotype *A* (in blue) carries one allele coding for specific rejection toward partners displaying phenotype *A* (in orange) and one allele coding for random mating (in grey). This butterfly will mate preferentially with individuals displaying phenotypes *B* and *C*, resulting in disassortative mating. 3. A butterfly displaying phenotype *A* (in blue) carries two alleles coding for specific rejection toward partners displaying phenotype *C* (in green). This butterfly will mate preferentially with individuals displaying phenotypes *A* and *B*. 4. A butterfly displaying phenotype *A* (in blue) carries one allele coding for specific rejection toward partners displaying phenotype *A* (in blue) and one allele coding for specific rejection toward partners displaying phenotype *C* (in green). This butterfly will mate preferentially with individuals displaying phenotype *B* resulting in disassortative mating. Cases 1 and 3 therefore lead to *self-acceptance*, while cases 2 and 4 lead to *self-avoidance*.

**Figure S8.**
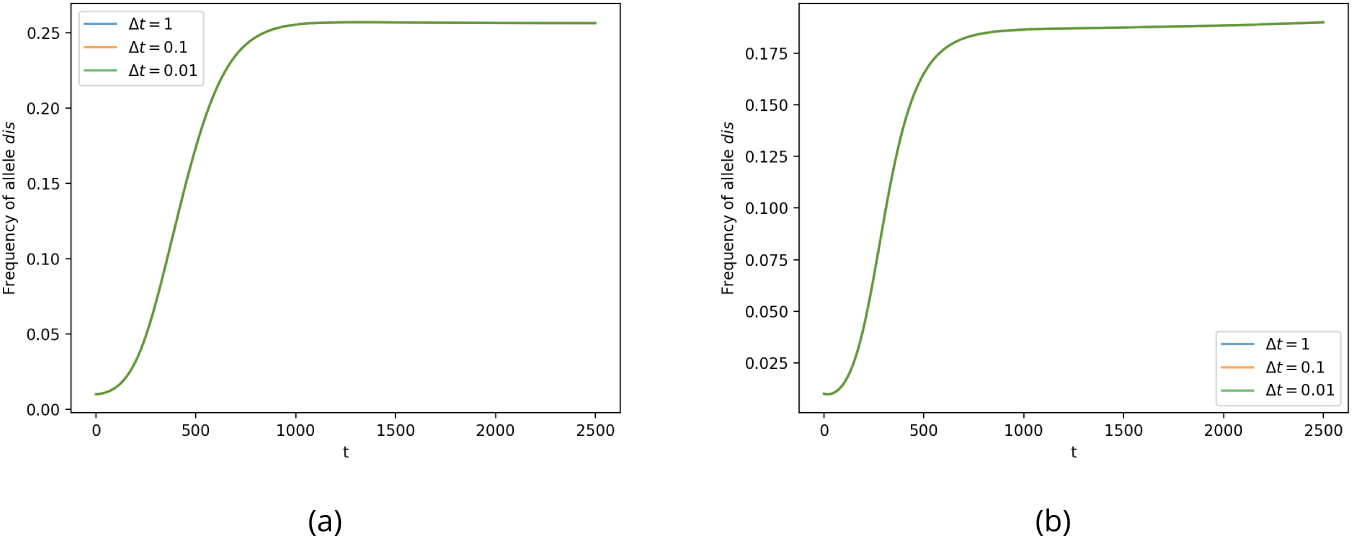
Evolution of the proportion of a mutant *dis* in the population immediately after its introduction, using simulations with three different time units (Δ*t* = 1 in blue, Δ*t* = 0.1 in orange or Δ*t* = 0.01 in green), under the *self-referencing* hypothesis (hyp. 1). All simulations give similar dynamics, assuming (a) *δ*_*a*_ = *δ*_*b*_ = 0.5, *δ*_*c*_ = 0 or (b) *δ*_*a*_ = *δ*_*b*_ = *δ*_*c*_ = 0.2, confirming that using discrete time simulations provides relevant estimations of the evolution of disassortative mating. Simulations are run during 2500 time steps and assuming, *r* = 1, *K* = 2000, 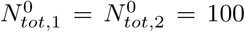, *λ* = 0.0002, *d*_*m*_ = 0.05, *d*_*n*−*m*_ = 0.15, *ρ* = 0, *mig* = 0.1, *δ* = 0.1 and *c*_*r*_ = 0.1.

**Figure S9.**
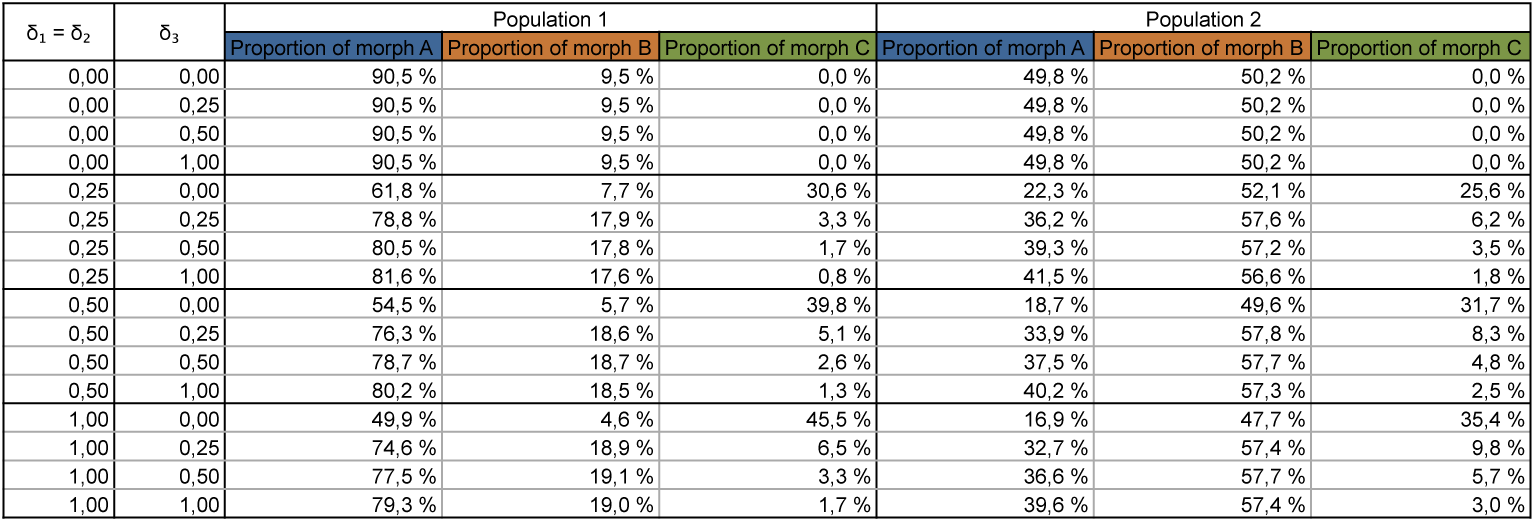
Influence of genetic load on color pattern polymorphism, assuming random mating. The proportions of phenotypes *A, B* and *C* in the populations living in patch 1 and 2 respectively at equilibrium depend on the different values of genetic load associated with the dominant allele *a* (*δ*_*a*_), intermediate-dominant allele *b* (*δ*_*b*_) and recessive allele *c* (*δ*_*c*_). Simulations are run assuming *r* = 1, *K* = 2000, 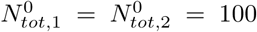, *λ* = 0.0002, *d*_*m*_ = 0.05, *d*_*n*−*m*_ = 0.15, *ρ* = 0, *mig* = 0.1, *δ* = 0.1 and *c*_*r*_ = 0.1.

**Figure S10.**
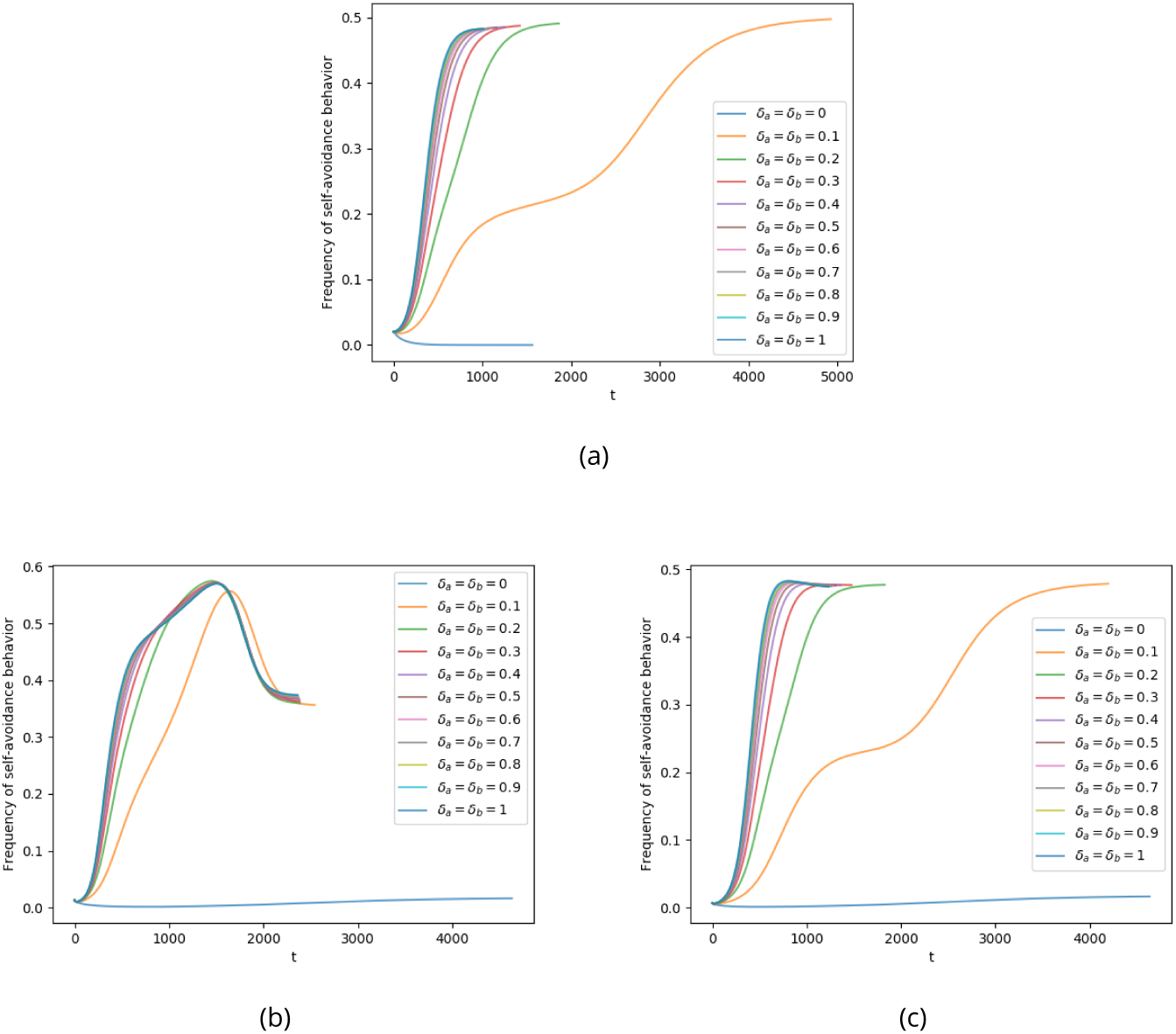
Frequency of *self-avoidance* behavior at the population level through time for different levels of genetic load, assuming (a) *self-referencing* (hyp. 1), (b) *attraction* rule (hyp. 2.a) or (c) *rejection* rule (hyp. 2.b) at the preference locus (*recognition/trait*). The evolution of the proportion of individuals displaying *self-avoidance P*_*s-av*_ after the introduction of preference alleles until equilibrium are shown for different values of genetic load *δ*_*a*_ and *δ*_*b*_. Simulations are run assuming *r* = 1, *K* = 2000, 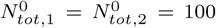, *λ* = 0.0002, *d*_*m*_ = 0.05, *d*_*n−m*_ = 0.15, *ρ* = 0, *mig* = 0.1, *δ*_*c*_ = 0, *δ* = 0.1 and *c*_*r*_ = 0.1.

**Figure S11.**
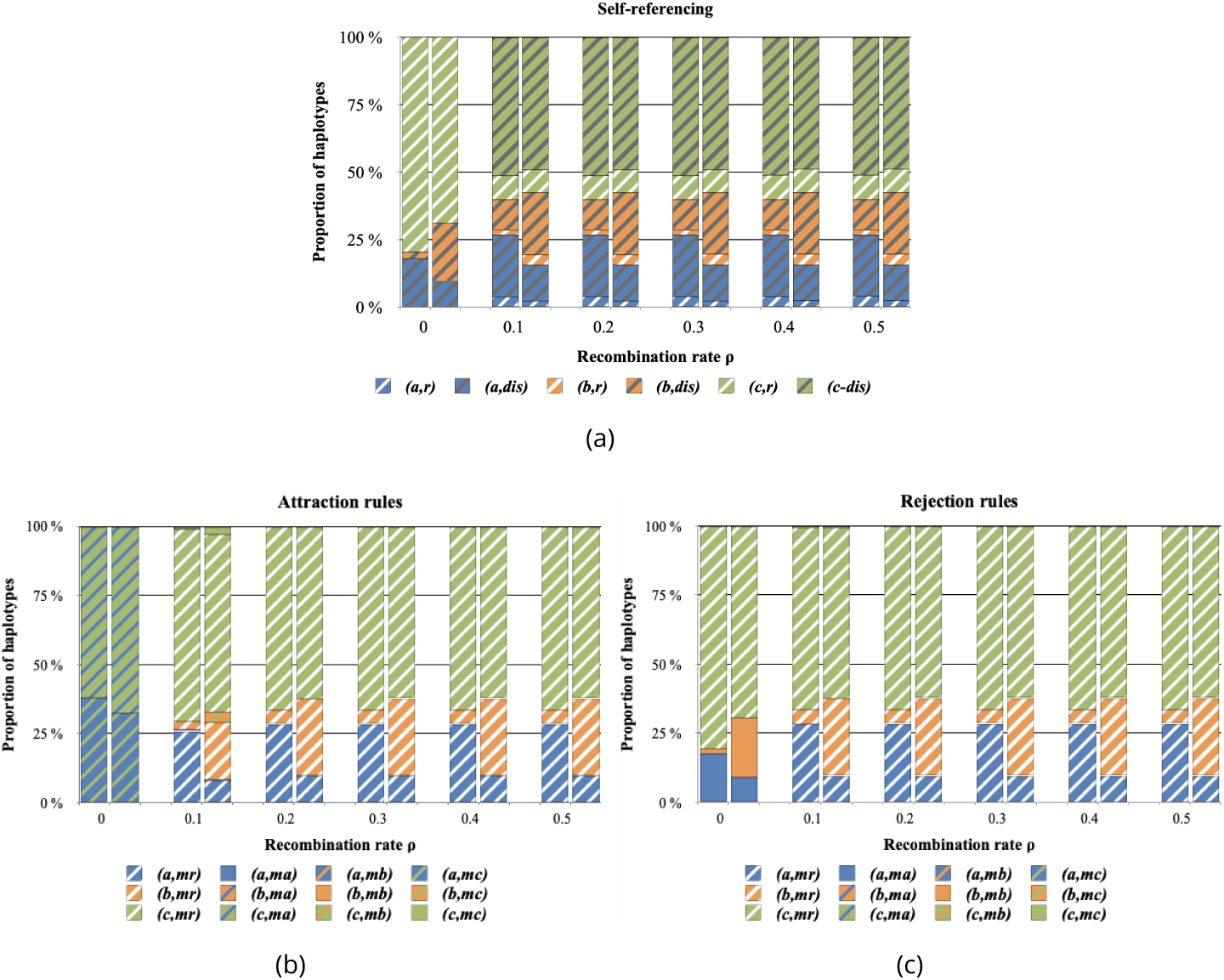
Influence of the recombination between color pattern and preference alleles on haplotype diversity, assuming (a) *self-referencing* (hyp. 1), (b) *attraction* rule (hyp. 2.a) or (c) *rejection* rule (hyp. 2.b) at the preference locus (*recognition/trait*). The proportion of haplotypes at equilibrium after the introduction of preference alleles in both patches are shown for different values of recombination rate *ρ* between the preference locus *M* and the color pattern locus *P*. For each value of recombination rate (*ρ*) the first and second bars represented haplotype proportions in the populations living in the patch 1 and 2 respectively. Simulations are run assuming *r* = 1, *K* = 2000, 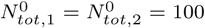, *λ* = 0.0002, *d*_*m*_ = 0.05, *d*_*n−m*_ = 0.15, *mig* = 0.1, *δ*_*a*_ = *δ*_*b*_ = 0.5, *δ*_*c*_ = 0, *δ* = 0.1 and *c*_*r*_ = 0.1.

